# Multi-omics reveal *Salmonella*-liberated dietary L-arabinose promotes expansion in superspreaders

**DOI:** 10.1101/2022.07.19.500711

**Authors:** Sarah Ruddle, Liliana M. Massis, Alyssa C. Cutter, Denise M. Monack

## Abstract

Our understanding of specific metabolites that drive host-pathogen interactions in the guts of superspreader hosts is incomplete. In a mouse model of chronic, asymptomatic *Salmonella enterica* serovar Typhimurium (*S*. Tm) infection, we performed untargeted metabolomics on the feces of mice and found superspreader hosts possess distinct metabolic signatures compared to non-superspreaders, including differential levels of L-arabinose. RNA-seq on *S*. Tm in superspreader fecal samples showed increased expression of the L-arabinose catabolism pathway *in vivo*. By combining bacterial genetics and diet manipulation, we demonstrate that diet-derived L-arabinose provides *S*. Tm a competitive advantage in the gut. We show that expansion of *S*. Tm in the gut requires a previously uncharacterized alpha-N-arabinofuranosidase that can liberate L-arabinose from dietary polysaccharides. Ultimately, we demonstrate that pathogen-liberated L-arabinose from the diet provides a competitive advantage to *S*. Tm *in vivo*. These findings propose L-arabinose as a critical driver of *S*. Tm expansion in the guts of superspreader hosts.

## INTRODUCTION

*Salmonella enterica* serovars are responsible for millions of infections worldwide. Disease states range from self-limiting gastroenteritis to invasive enteric fever, the latter of which is caused by human-adapted *S*. *enterica* serovar Typhi (*S*. Typhi) (Parry *et al*., 2002). In mouse oral infection models, *S*. *enterica* serovar Typhimurium (*S*. Tm) robustly colonizes the cecum and colon (Lam and Monack, 2014). From these sites *S*. Tm is shed via the feces into the environment, where it can infect naïve hosts (Crum Cianflone, 2009). Transmission efficiency has been linked to increased levels of fecal bacteria, with high-shedding hosts (referred to as superspreaders) responsible for most transmission events (Gopinath et al., 2012). A subset of infected individuals become carriers and persistently shed *Salmonella* in their feces, thereby functioning as a reservoir for the pathogen. The most famous superspreader, Mary Mallon, is remembered by the moniker Typhoid Mary for the dozens whom she infected with *S*. Typhi as a cook at the turn-of-the-century (Soper, 1939; McGovern and Leavitt, 1997). For transmission events to occur, *Salmonella* utilizes several key virulence factors such as *Salmonella* Pathogenicity Islands (SPI-1), SPI-2, and SPI-6, which encode secretion systems that deliver effector proteins into host cells and competing Enterobacteriaceae (Lam and Monack, 2014; Sana et al., 2016) to establish itself in the gastrointestinal tract and replicate to high levels while concurrently competing against commensal microbiota for resources. Although microbiome-*Salmonella* interactions have been shown to influence *Salmonella* shedding levels (Rogers, Tsolis and Bäumler, 2021), specific features that distinguish superspreaders from non-superspreaders remain unknown.

The complex microbial community occupying the gastrointestinal tract plays a crucial role in preventing pathogen invasion through direct and indirect mechanisms, referred to as colonization resistance (Sorbara and Pamer, 2019). We and others have demonstrated that the commensal microbiota provide colonization resistance against *S*. Tm infection, as oral antibiotic treatment enhances pathogen expansion and fecal shedding (Barthel *et al*., 2003; Lawley *et al*., 2008). Mechanisms of colonization resistance against intestinal *S*. Tm infection have mainly been studied in colitis models of infection, where mice are pre-treated with antibiotics resulting in high levels of inflammation (Barthel *et al*., 2003). Multiple studies have also used this colitis model to shed light into metabolic pathways that *S*. Tm uses to expand in the guts of antibiotic-treated mice (Rogers et al., 2021). For example, S. Tm takes advantage of nitrate generated by the host immune response by using it as an electron acceptor to fuel anaerobic respiration (Barrett and Riggs, 1982), which allows *S*. Tm to outgrow the resident microbiota (Lopez *et al*., 2012; McLaughlin *et al*., 2019). The inflamed gut is a unique niche in which *S*. Tm alters its metabolism and begins to utilize carbon sources that require respiration such as ethanolamine, fructose-asparagine, 1,2-propanediol, and propionate (Thiennimitr *et al*., 2011; Ali *et al*., 2014; Faber *et al*., 2017; Shelton *et al*., 2022). Although important discoveries were made, disruption of the resident microbial community in this antibiotic-treated mouse model limits the generalizability of mechanistic findings to clinical infections in humans, in which some individuals become chronic asymptomatic superspreaders. Thus, in this study, we use a mouse model of oral *S*. Tm infection in which mice are not pre-treated with an antibiotic to uncover mechanisms of *Salmonella* colonization that promote the emergence of superspreaders.

In a conventional mouse model, *Salmonella* must directly compete with commensal microbiota for a nutritional niche (Sommer *et al*., 2017). Diets contain complex carbohydrates, the main component of dietary fiber, and are the primary metabolic input for both commensal and pathogenic bacteria (Sonnenburg and Sonnenburg, 2014). The ability of commensal microbes to process complex carbohydrates is critical for their colonization of the gut. Commensal bacteria process complex carbohydrates through glycoside hydrolases that cleave specific glycosidic linkages to liberate oligosaccharides or monosaccharides for catabolism (Wardman *et al*., 2022a). Some bacteria have evolved to sequester free saccharides from competing microbes through mechanisms of rapid import systems (Martens et al., 2009); others can work in unison to break down complex carbohydrates in so-called cross-feeding events (Martens *et al*., 2014). *Salmonella* can participate in cross-feeding by catabolizing commensal-liberated fucose and sialic acid from host-derived mucus for its colonization in colitis models (Ng *et al*., 2013). Although these studies have advanced our understanding of dietary carbohydrate processing by commensals, little is known about how pathogens like *Salmonella* utilize complex sugars during infection and how these metabolic processes affect superspreaders.

The metabolic networks of host, microbiome, and pathogen are interconnected during infection, and a deeper understanding of the metabolic signatures that underlie superspreaders may provide insights relevant for infection control. In this study, we performed untargeted metabolomic analysis in superspreader and non-superspreader mice to identify a metabolic state that is unique to superspreaders. We narrowed our focus to pathogen-specific factors by analyzing the transcriptional profile of *S. Tm* from superspreader feces and identified L-arabinose metabolism as a critical pathway for *Salmonella* expansion in the gut. We discovered that exogenous supplementation of dietary L-arabinose promotes the rapid emergence of superspreaders. We demonstrate that L-arabinose catabolism confers a strong competitive advantage for *Salmonella* in the gut, which was independent of the microbiome. Finally, we identified a previously uncharacterized alpha-n-arabinofuranosidase in *Salmonella* that plays a key role during colonization of the gut. Collectively, these findings uncover a metabolite-dependent mechanism of pathogen expansion within a mouse model of chronic *Salmonella* infection.

## RESULTS

### *Salmonella* superspreaders have a distinct metabolic signature

The molecular mechanisms that impact *Salmonella* expansion in the distal guts of chronically infected mice are incompletely understood. To gain insight into mechanisms of pathogen expansion, ninety-five 129X1/SvJ mice were intragastrically inoculated with the wild-type (WT) *S*. Tm strain SL1344 and bacterial loads were measured in the feces at 7-, 14-, and 21-days post infection (DPI) (Figures 1A, S1A). As previously published, mice shed a range of *S*. Tm in the feces over time with approximately 20% of mice shedding > 10^8^ colony forming units (CFU)/g feces, which we define as superspreaders (Gopinath, Carden, & Monack, 2012) (Figure 1B). In addition, spleen, liver, mesenteric lymph nodes, cecal contents, and feces were collected 28 DPI to quantify *S*. Tm levels at these different infection sites for comparison to fecal shedding levels. Although the pathogen burden in the feces varied between the mice from the level of detection to 10^9^ CFU/g feces, there were no significant differences in the levels of *S*. Tm in the systemic tissues (Figure 1C). The heterogeneity of *S*. Tm levels was restricted to the distal gut (Gopinath et al., 2014), suggesting that distal gut-specific metabolites may affect pathogen expansion.

**Figure 1.**
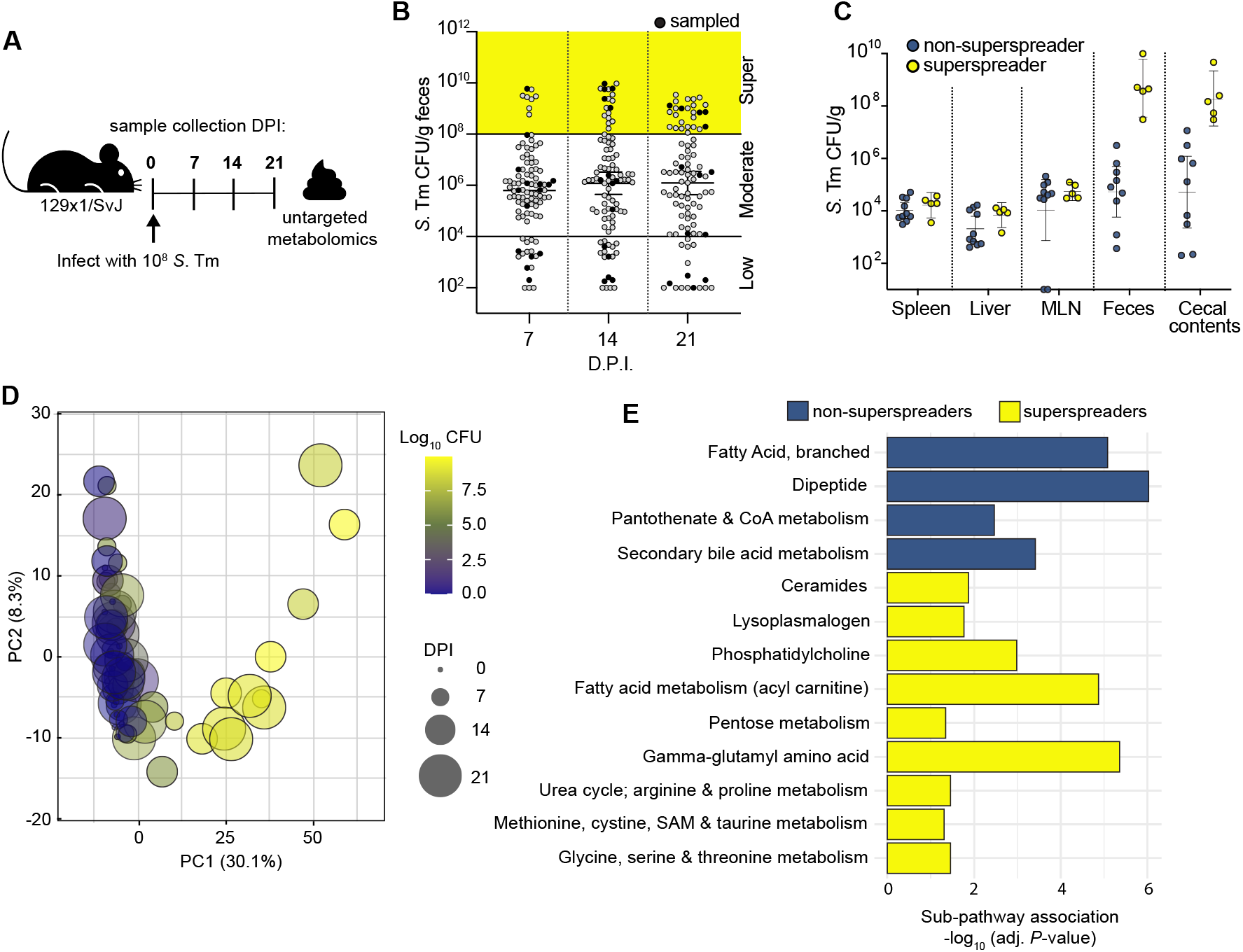
*S*. Tm superspreaders share a distinct metabolic landscape. A) Infection schematic for metabolomic profiling. 129X1/SvJ mice infected with 10^8^ CFU *S*. Tm for untargeted fecal metabolomics. Mice were the sampled at the indicated timepoints: 0,7,14, and 21 days post-infection. B) *S*. Tm CFU/g feces from 129X1/SvJ 7,14, and 21 days post-infection (N=95). Black circles indicate mice selected from each shedding stratum (low, moderate, supershedding; n=5) for fecal metabolite profiling. C) *S*. Tm burden across organs of 15 mice selected for fecal metabolomics day 28 post-infection. N=5 mice per shedding strata group. D) Principal component analysis of fecal metabolomes from 20 mice, over 4 timepoints, colored by CFU, and sized by day post infection (0-21 DPI). E) Differential pathway enrichment of non-superspreaders and superspreaders at 14 DPI with multiple hypothesis correction (likelihood ratio test with Benjamini & Hochberg correction). See also Figure S1.

To better understand how the metabolic landscape of the intestine changes during *S*. Tm infection, we performed unbiased, semi-quantitative profiling on extracellular metabolites in the gut lumen over time. Specifically, fecal samples from mice shedding low (10^2^ – 10^4^ CFU/g), moderate (10^4^ - 10^8^ CFU/g), or superspreader (>10^8^ CFU/g) levels of pathogen (n=5 per shedding group; Figure 1A) were collected and subjected to ultra-high-performance liquid chromatography-tandem mass-spectrometry (UHPLC/MS). The relative abundances of 797 metabolites in the feces were measured in infected mice over time and compared to mock-treated mice (Metabolon, USA). The analysis of the longitudinal samples identified a distinct metabolic signature that developed in superspreaders over time. Principal component analysis (PCA) of the full dataset showed the effect of *S*. Tm burden (log CFU) on the metabolome was largely captured by PC1 (Figure 1D), with the first two PCs accounting for 38.4% of the variation in the dataset. PCA showed partitioning by shedding stratum (Figure 1D), where superspreaders clustered together and away from low and moderate shedders, indicating that these superspreaders share a distinct metabolome compared to the low and moderate shedders.

Pathway enrichment analysis identified metabolic pathways with significant differential enrichment in superspreader and non-superspreader mice throughout the infection (Figure1E and Figures S1B-1C, likelihood ratio test with Benjamini & Hochberg correction). Acyl-carnitine, pentose, gamma-glutamyl amino acid, and phosphatidyl choline metabolism are several of the pathways enriched in superspreader versus non-superspreader feces on 14 DPI (Figure 1E). Enriched pathways of non-superspreader feces collected on 14 DPI include secondary bile acid, dipeptide, and branched fatty acid metabolism pathways (Figure 1E). Comparing pathway enrichment across timepoints highlights the metabolic flux occurring over the time course of infection. For example, acyl-choline and tocopheral metabolism are enriched in superspreader mice versus non-superspreader mice only at 7 DPI (Figure S1B). In contrast, metabolic pathways like ceramides associate with superspreaders at 14, and 21 DPI (Fig 1E and Figure S1B-1C.). Together, these findings suggest that the metabolic landscape of superspreader mice is distinct and may reflect alterations in luminal nutrient availability during *S*. Tm infection.

### *S. Tm* gene expression in guts of superspreader mice

We performed metatranscriptomic analysis to prioritize *S*. Tm phenotypes that support expansion in the distinct metabolic environment of superspreaders. Metatranscriptomics quantifies mRNA across the gastrointestinal microbial community and can demonstrate which genes are differentially expressed between contexts. Mice were infected with wild-type *S*. Tm strain SL1344 and fecal shedding levels were monitored over time (Figure 2A). After 7,14, and 21 DPI, mice were euthanized, and cecal contents and feces were collected for total RNA extraction (including host, commensal microbes, and *S*. Tm RNA). Three representative superspreader mice at each timepoint were used for bulk fecal RNA-seq (Figure 2A). In addition to the *in vivo* samples, RNA was extracted from *S*. Tm grown in LB cultures, grown both aerobically and anaerobically to mid-log, log, and stationary stage. PCA analysis comparing *S*. Tm transcriptomes of all *in vitro* versus all *in vivo* conditions showed that the *in vivo* data separated from *in vitro* samples along PC1 (Figure 2SA). We found 1755 genes that were differentially expressed between *in vivo* (cecum and feces) and *in vitro* S. Tm (adjusted P<0.05, Likelihood ratio test with Benjamini & Hochberg correction).

**Figure 2.**
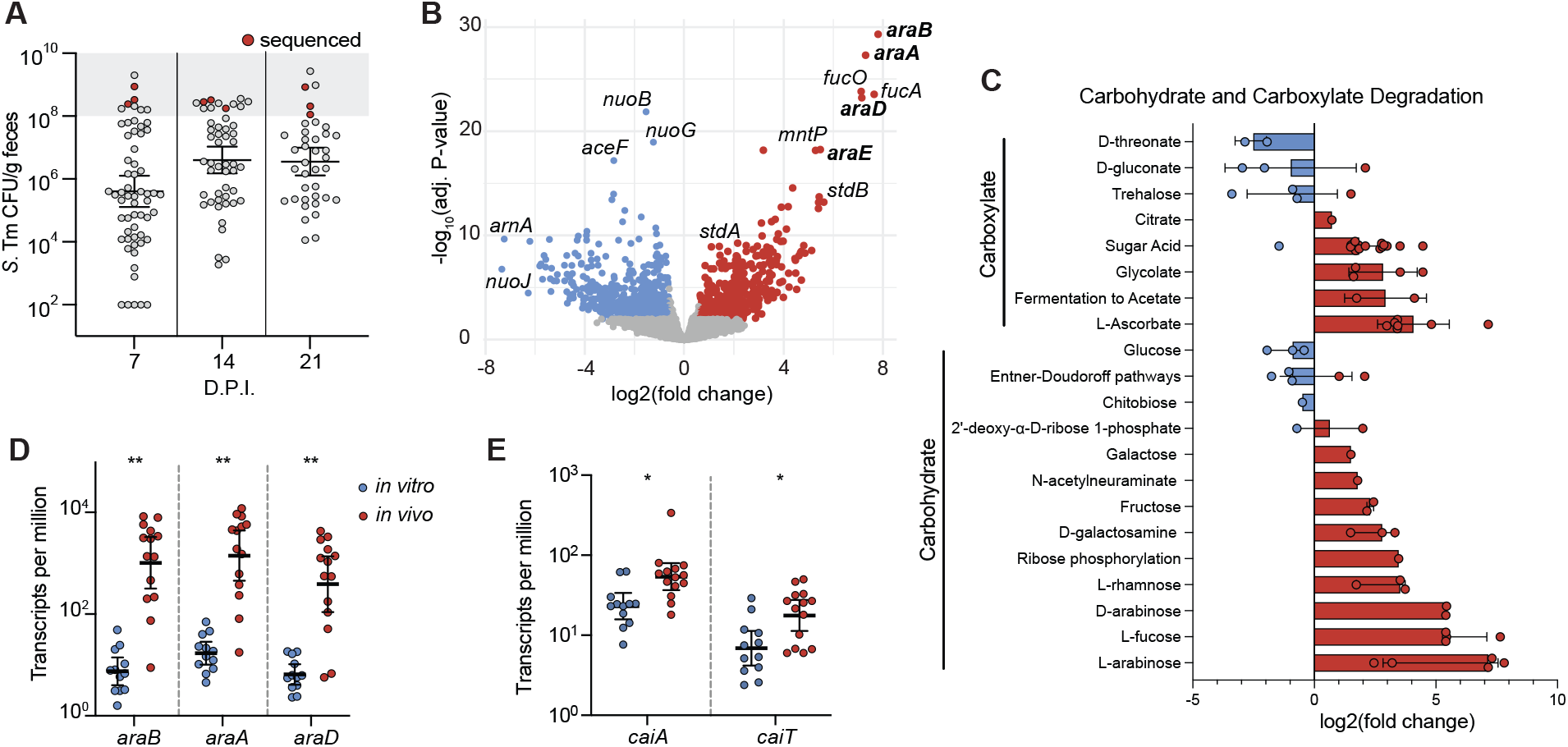
*In vivo* fecal transcriptomics of *S*. Tm superspreaders reveal changes in carbohydrate utilization. A) 129X1/SvJ mice infected with 10^8^ CFU WT *S*. Tm for *in vivo* RNA seq. *S*. Tm shedding levels over time, red data points indicate superspreader mice selected for metatranscriptomic analysis of feces on 7, 14,and 21days post-infection. N=3 at each timepoint. B) Significant differentially expressed genes *in vitro* (blue) and *in vivo* (red) and not significantly enriched in grey, -log_10_ adjusted p-value and log2 fold change. C) Log2 fold change of differentially enriched metabolites (adjusted p-value < 0.05; Benjamini & Hochberg correction) *in vitro* (blue) and *in vivo* (red). Each data point represents single metabolites within each sub pathway (y-axis). Sub pathways are annotated with corresponding major pathways (brackets along y-axis). D) Transcripts per million of *araB, araA, and araD in vitro* (blue) and *in vivo* (red). ** p < 0.01 T test with Welch’s correction E) Transcripts per million of *caiA and caiT in vitro* (blue) and *in vivo* (red). * p < 0.05 T test with Welch’s correction See also Figure S2.

Genes involved in several metabolic pathways were enriched in *S*. Tm from superspreader mice compared to *in vitro* samples (Figure 2B). Genes that encode known virulence factors, such as *ssaV, sifA, pipB2*, and *sseI*, were expressed at lower levels in the gut compared to *in vitro* growth conditions, which is consistent with the expression of these virulence factors being important for *S*. Tm growth and survival in tissues rather than growth in the gut lumen (Diard *et al*., 2013; Hockenberry *et al*., 2021). In contrast, genes previously demonstrated to impact *Salmonella* colonization of the gut, such as fucose metabolism (*fucA, fucO, fucK*) (Deatherage Kaiser *et al*., 2013; Ng *et al*., 2013) and fimbriae genes (*stdA, stdB*) (Chessa *et al*., 2008; Suwandi *et al*., 2019) were expressed at higher levels in the guts compared to *in vitro* growth conditions (Figure 2B). We also plotted the log2 fold change of all significantly enriched genes by their functional pathway to visualize an overview of transcriptional changes *S*. Tm undergoes in superspreader hosts versus *in vitro* growth conditions (Figure S2B). We observed overlapping enriched pathways from the transcriptomics and the metabolomics. For example, gene transcripts from the carbohydrate and carboxylate degradation pathways were enriched *in vivo* (Figure S2B), and changes in pentose metabolites were enriched in superspreaders (Figure 1E). To look closer at carbohydrate catabolism dynamics in *S*. Tm, we plotted the log2 fold change of genes involved in the carbohydrate and carboxylate degradation pathways and found L-arabinose degradation to be the most enriched pathway *in vivo* (Figure 2C). Interestingly, glucose degradation gene transcripts were lower *in vivo* than *in vitro*, which agree with the cross regulation of L-arabinose and glucose utilization in *S*. Tm (Chakravorty, 1964). Transcripts from the L-arabinose utilization operon (*araB, araA, araC*) isolated from *S*. Tm *in vivo* were ~100-fold higher compared to *in vitro* (Figure 2D).

In addition to L-arabinose catabolism, levels of the carnitine catabolism gene transcripts (*caiA, caiT*) were enriched in *S*. Tm isolated from superspreader guts compared to *S*. Tm grown *in vitro* (Figures 2E), and metabolomics data indicated that superspreaders have a unique carnitine metabolic profile, since the acyl-carnitine pathway was enriched in superspreaders (Figure 1E). Together, these data suggest that *S*. Tm catabolism of L-arabinose and carnitine may play roles in colonization of superspreader guts.

### L-arabinose and carnitine metabolism changes in superspreader mice

The increased expression of the genes involved in L-arabinose and carnitine metabolism from *S*. Tm isolated from superspreader mice led us to focus on understanding the roles of these pathways *in vivo*. We observed significant differences in the pentose metabolism and acyl carnitine sub-pathways in superspreaders compared to non-superspreaders (Figure 1E). The scaled level of L-arabinose decreased in superspreader mice over time, yet remained constant across uninfected, low, and moderate shedders (Figure 3A). We confirmed the levels of L-arabinose by quantifying the L-arabinose/g feces of uninfected mice and superspreader mice at 14 DPI through targeted LC/MS. Like the scaled metabolite levels, there were significantly lower levels of L-arabinose in the feces of superspreader mice compared to uninfected mice (Figure 3B). These data led us to hypothesize that *S*. Tm consumes L-arabinose in the gut lumen of superspreader mice. Although, L-arabinose metabolism in *Enterobacteriaceae* has been relatively well characterized (Ammar *et al*., 2018; Schleif, 2021), the role of L-arabinose metabolism during infection has not been studied in detail. In *Salmonella enterica* serovars, the primary L-arabinose permease is AraE (López-Garrido *et al*., 2015). Intracellularly, the ribulokinase, AraB, and the L-arabinose isomerase, AraA, produce L-ribulose-5-phosphate from L-arabinose. AraD, an L-ribulose-5-phosphate 4-epimerase, then forms L-xylulose-5-phosphate. L-xylulose-5-phosphate is further processed in known down-stream pathways ultimately feeding into the Pentose Phosphate Pathway or the TCA cycle (Mayer and Boos, 2005) (Figure 3C). Most metabolites within the acyl-carnitine pathway increased in superspreader mice over time, yet remained constant across low, and moderate shedders (Figure 3D). Upon import into the cell through CaiT, the L-carnitine:gamma-butyrobetaine antiporter, *S*. Tm can degrade L-carnitine as outlined in Figure 3E (Meadows and Wargo, 2015). The changes in both the metabolites and gene expression of carnitine-related metabolites led us to question if *S*. Tm could benefit from the surplus of carnitine derivatives in the gut environment.

**Figure 3.**
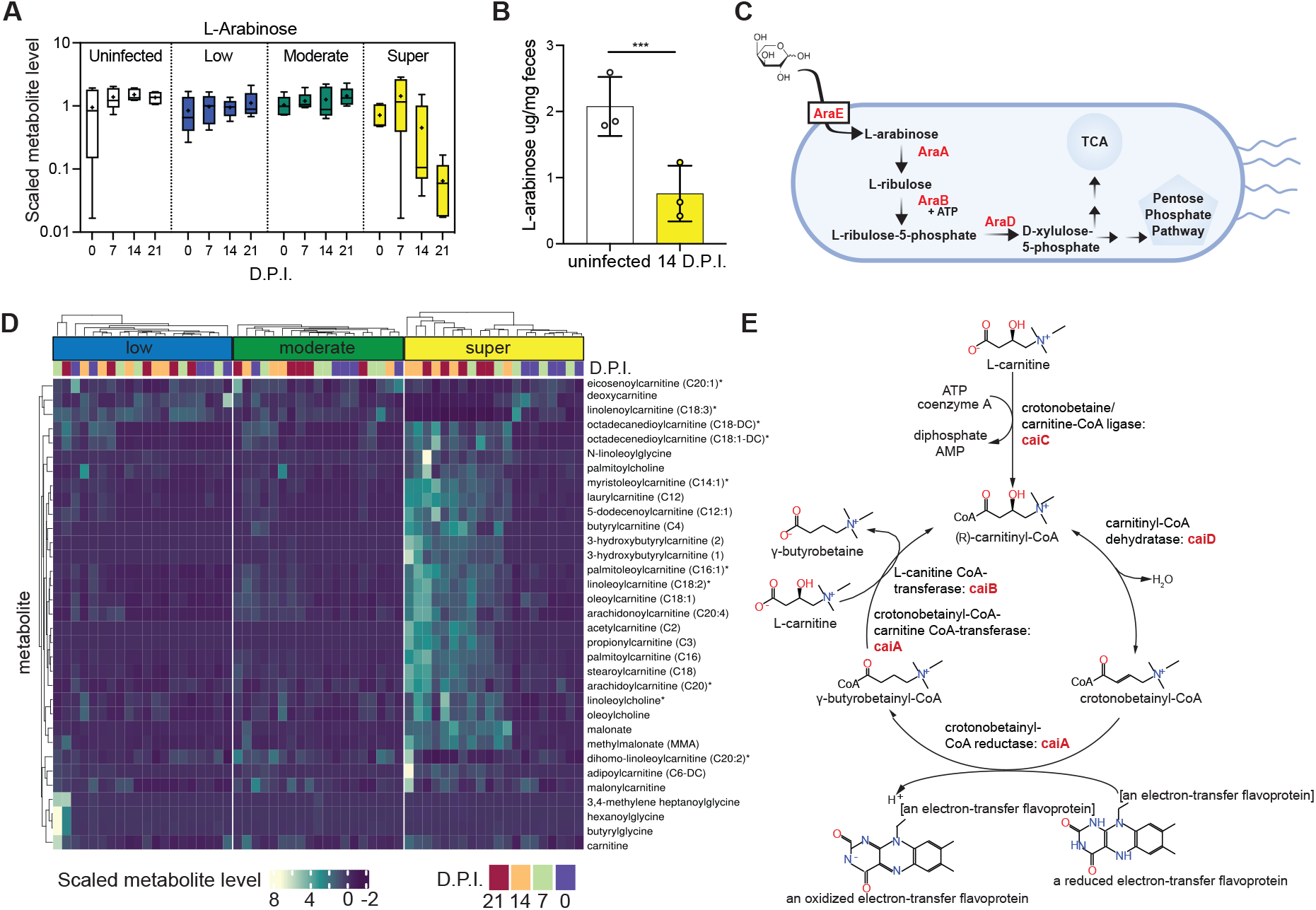
L-arabinose metabolism is important for *S*. Tm expansion in superspreader mice. A) Scaled-metabolite level of L-arabinose detected in stool of uninfected and infected mice colonized across shedding strata. B) L-arabinose levels detected in uninfected mice and superspreader mice feces (collected 14 DPI) by targeted LC/MS. T test *** p < 0.001 C) Simplified schematic of L-arabinose catabolism in *Salmonella enterica* D) Heat map showing scaled metabolite levels of metabolites from carnitine metabolism pathway for all shedding strata and four timepoints (0-21 DPI) E) Schematic of L-carnitine catabolism in *S*. Tm

### L-arabinose catabolism confers competitive advantage to *S*. Tm in the gut lumen

The overlap of our metabolomic and transcriptomic data in superspreaders led us to hypothesize that *S*. Tm exploits L-arabinose and/or L-carnitine to expand in the distal gut. To interrogate the importance of these pathways, we provided 1% exogenous L-arabinose or 1% L-carnitine in the drinking water of mice at the start of infection and maintained mice on the treated water throughout the course of infection. Strikingly, *S*. Tm expanded rapidly in the guts of mice provided exogenous L-arabinose, where the average shedding levels in mice provided exogenous L-arabinose were >100 fold higher than control mice by 3 DPI (Figure 4A). In contrast, mice given exogenous L-carnitine shed similar levels of *S*. Tm compared to mice given control (standard) water throughout the course of infection (Figure 4B). The dramatic expansion of *S*. Tm in mice supplemented with L-arabinose led us to question the indirect effect exogenous L-arabinose could be having in the mice. Inflammation has been well-established to facilitate *S*. Tm expansion in the gut by various mechanisms that include use of an array of exogenous respiratory electron acceptors, consumption of fermentation products, or oxidized sugars (Rivera-Chávez and Bäumler, 2015). To determine if exogenous L-arabinose was creating an inflammatory environment for *S*. Tm to exploit, we profiled fecal levels of inflammatory markers such as lipocalin-2 and proinflammatory cytokine mRNA expression in the presence or absence of exogenous L-arabinose from uninfected mice and mice infected with *S*. Tm. There was no significant difference in lipocalin-2 levels at 4 DPI (Figure S3A) in the feces or mRNA transcripts of inflammatory cytokine interferon gamma (Figure S3B-C) at 5 DPI in the colon or spleen. Together these data suggest *S. Tm* uses L-arabinose to expand in the gastrointestinal tract, independent of inflammation. Ruling out indirect inflammation brought on by increased exogenous L-arabinose led us to hypothesize arabinose is an important carbon source for *S*. Tm in the gut.

**Figure 4.**
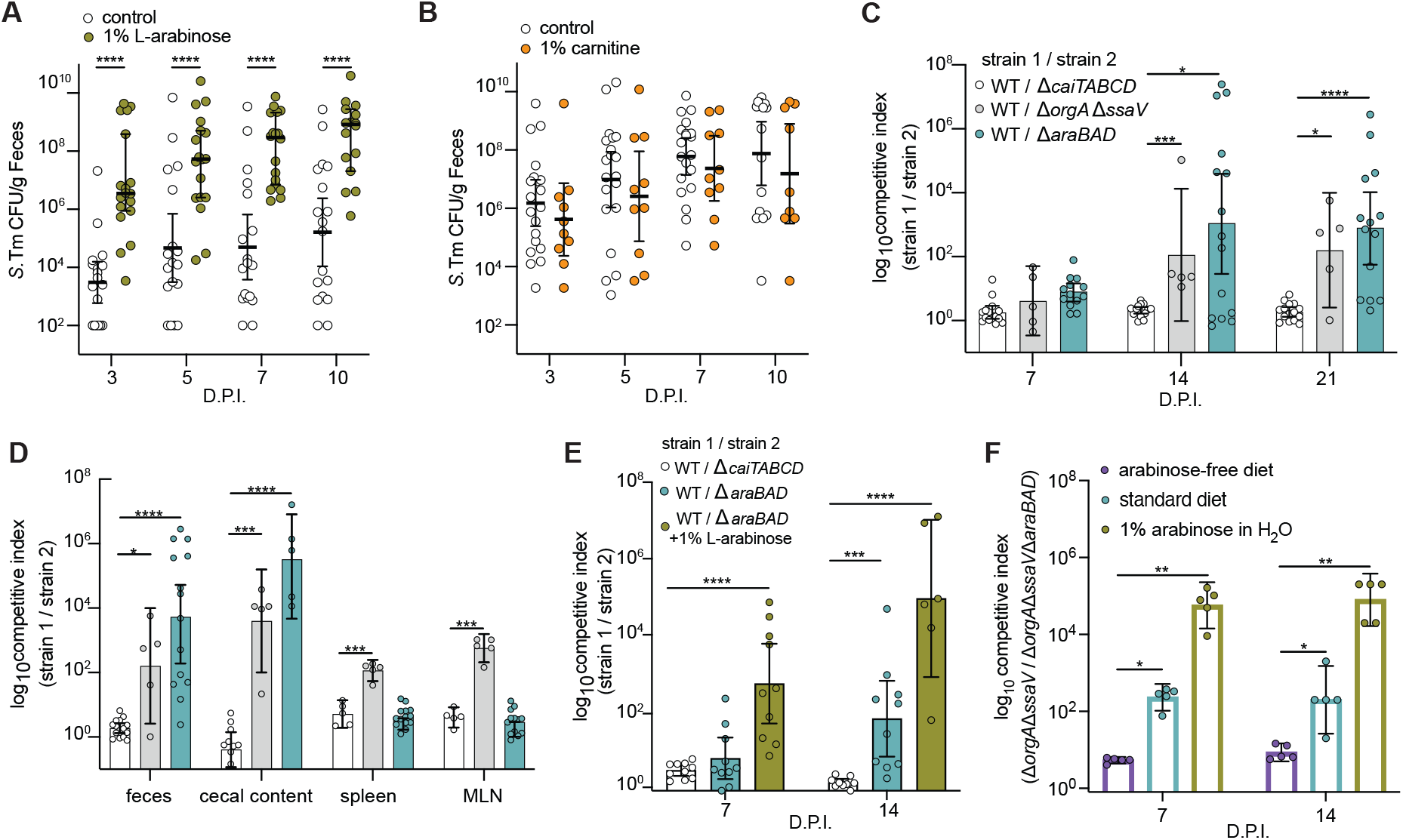
L-arabinose utilization confers a competitive advantage to S. Tm *in vivo*. A) 129X1/SvJ mice infected with WT *S*. Tm and *S*. Tm CFU/g feces quantified over 10-days. White dots indicate control mice (n=18) and green dots represent mice provided 1% exogenous L-arabinose through drinking water (n=18). *** p < 0.001, **** p < 0.0001 Multiple Mann-Whitney tests B) 129X1/SvJ mice infected with WT *S*. Tm and *S*. Tm CFU/g feces quantified over 10-days. White dots indicate control mice (n=20) and orange dots represent mice provided 1% exogenous L-carnitine through drinking water (n=10). No significant difference between groups by Multiple Mann-Whitney tests C) 129X1/SvJ mice were inoculated with an equal mixture of the indicated strains. The competitive index of the two strains in the feces was determined at 3-, 7-, 14-, and 21-days post – infection. D) 129X1/SvJ mice (from fig. 4A) were euthanized 21 DPI for organ collection. Spleen, mesenteric lymph nodes, cecal content, and feces was homogenized and plated for CFU and competitive index. E) 129X1/SvJ mice were inoculated with an equal mixture of the indicated strains. One group of mice were provided 1% L-arabinose drinking water (green). The competitive index in the stool was determined on 7- and 14-days post-infection. N=10 mice per group F) Germ free mice were given the indicated dietary conditions one day prior to infection. Mice were infected with an equal mixture of *DorgAssaV and DorgAssaVDaraBAD*. Competitive index was determined at 7 and 21 days post-infection. N=5 per group. Each dot represents data from one animal (biological replicate). Bars represent +-SEM. *p < 0.05, ** p < 0.01, *** p < 0.001, **** p < 0.0001. Mann – Whitney U test unless otherwise stated. See also Figure S3 and S4.

To test if L-arabinose utilization or carnitine degradation pathways confer a fitness advantage for *S*. Tm in the gut, we infected mice with an equal mixture of WT *S*. Tm and isogenic mutants and calculated a competitive index (CI) at multiple timepoints throughout the course of infection. Groups of 129X1/SvJ mice were orally inoculated with an equal mixture of the *S*. Tm WT strain and either an L-arabinose catabolism mutant strain, Δ*araBAD::Kan^R^* or a carnitine catabolism mutant strain, *DcaiTABCD::Kan^R^*. As a positive control, mice were orally infected with an equal mixture of WT *S*. Tm and a strain that is deficient for critical components of the Type Three Secretion Systems (T3SS) in *S*. Tm, *DorgAΔssaV::Kan^R^ S*. Tm, and is attenuated for colonization of conventional mice (Trevor D. Lawley *et al*., 2008). As expected, WT *S*. Tm outcompeted the *DorgAssaV::Kan^R^ S*. Tm, with CI of ~100 at 14 DPI (Figure 4C). The WT and *DcaiTABCD::Kan^R^* S. Tm strains were equally fit during the 21 day infection (Figure 4C). Since the L-carnitine-deficient mutant did not have a defect in the guts of mice, we utilized this mutant as a control for the chromosomal insertion of the kanamycin resistance gene in subsequent competition experiments. We found that the Δ*araBAD::Kan^R^ S*. Tm strain suffered a competitive disadvantage after 14 DPI compared to WT *S*. Tm, with a CI of >1000 in the guts of mice (Figure 4C), which was even higher than the fitness defect observed with the control T3SS-deficient strain. Collectively, our data indicate that L-arabinose metabolism provides a competitive advantage to *S*. Tm in the gut.

To determine whether *S*. Tm utilization of L-arabinose confers a competitive advantage in other tissues, we measured *S*. Tm levels in cecal content, mesenteric lymph nodes (MLN), and spleen at 21 DPI. The WT *S*. Tm strain outcompeted the Δ*araBAD::Kan^R^ S*. Tm strain in the cecal contents (Figure 4D). In contrast, the WT and Δ*araBAD::Kan^R^ S*. Tm strains were equally fit in the MLN and spleen (Figure 4D), indicating that L-arabinose metabolism confers a competitive advantage to *S*. Tm specifically in the gastrointestinal tract.

These results led us to hypothesize that dietary L-arabinose supplementation would lead to greater competition between the WT and the L-arabinose catabolism-deficient *S*. Tm mutant strains. To test this hypothesis, two groups of 129×1/SvJ mice were infected with a 1:1 mixture of WT and Δ*araBAD::Kan^R^ S*. Tm strains. One of these two groups received 1% exogenous L-arabinose in the drinking water at the time of infection and throughout the infection. We found that the kinetics by which the WT *S*. Tm strain outcompeted the Δ*araBAD::Kan^R^* strain were significantly faster in mice given L-arabinose in their water (Figure 4E). Specifically, the WT strain outcompeted the Δ*araBAD::Kan^R^ S*. Tm strain as early as 7 DPI with a CI~1,000 and by two weeks, the CI was ~10,000 (Figure 4E). Together, these results are consistent with a model in which L-arabinose is a carbon source available to *S. Tm* and that its catabolism through *araBAD* contributes to gut colonization and expansion.

In parallel we tested if exogenous L-carnitine would lead to a competitive advantage for WT *S*. Tm compared to Δ*caiTABCD::Kan^R^* S. Tm. Mice were given standard water or 1% L-carnitine water throughout the course of infection. To ensure mice were ingesting the 1% L-carnitine water, we gavaged additional groups of mice daily with 200μl of standard water or 1% L-carnitine water. There was no competitive advantage to WT *S*. Tm in any of the tested conditions (Supplemental Figure 4A). Together these data suggest *S*. Tm does not directly benefit from L-carnitine degradation during chronic mouse infection. With a competitive index of ~1, Δ*caiTABCD::Kan^R^* S. Tm continues to serve as a control for the chromosomal insertion of the kanamycin resistance gene. Moving forward, we chose to continue probing mechanisms of L-arabinose metabolism during *S*. Tm infection.

### Competitive advantage of *S*. Tm L-arabinose catabolism in the gut is not dependent on commensal bacteria

Given the critical role that L-arabinose utilization plays during *S. Tm* colonization of the gut, we next sought to understand how *S*. Tm is acquiring L-arabinose in the gut lumen of superspreader mice. In mice, L-arabinose is found within plant polysaccharides of fibrous dietary components and requires microbe-specific enzymes (glycoside hydrolases) to liberate free L-arabinose (Martens *et al*., 2011). To determine if the microbiota was responsible for liberating free L-arabinose in the gut lumen for subsequent use by *S*. Tm during infection, we performed a series of CI experiments in germ-free mice. One day prior to infection, germ-free mice were given either a standard diet and standard water, an arabinose-free diet and standard water, or standard diet and water containing 1% L-arabinose. However, WT *S. Tm* is lethal to germ-free mice (Collins and Carter, 1978), and both T3SS must be functionally inactive for *S*. Tm to colonize the mouse gut while maintaining host viability (Stecher *et al*., 2005). Thus, to assess *S. Tm* fitness and metabolism in the absence of other commensal microbes, germ-free mice were infected orally with a 1:1 mixture of Δ*orgAssaV::Strep^R^ and ΔorgAssaVΔaraBAD::Kan^R^ S*. Tm strains. In mice fed an arabinose-free diet, the Δ*orgAssaV::Strep^R^* strain had no competitive advantage compared to the L-arabinose-utilization mutant strain, Δ*orgAssaVΔaraBAD::Kan^R^* (Figure 4F). In contrast, the Δ*orgAssaV::Strep^R^* strain was significantly more fit than the strain lacking *araBAD* in mice given standard chow, which contains L-arabinose in the form of crude plant material (dietary fiber), and standard water (Figure 4F). Furthermore, we found that the Δ*orgAssaV::Strep^R^* strain outcompeted the Δ*orgAssaVΔaraBAD::Kan^R^* strain more rapidly and significantly in mice fed a standard diet and 1% L-arabinose water (Figure 4F). We also confirmed through LC/MS, that L-arabinose levels decrease during *S*. Tm colonization in the gnotobiotic (ex-germ-free), similar to our findings in the conventional 129X1/SvJ mouse model (Supplemental Figure 4B). Together these data reveal that *S*. Tm has a competitive advantage when able to catabolize L-arabinose, even in the absence of a commensal microbiota.

### *S*. Tm arabinofuranosidase confers competitive advantage to *S*. Tm in the gut lumen

Since the increased fitness of *S*. Tm conferred by L-arabinose catabolism is not dependent on the microbiome, we hypothesized that *S*. Tm has the functional capacity to liberate free L-arabinose from plant polysaccharides for its own use *in vivo*. To test this hypothesis, we first performed a BLAST search to identify potential arabinose-liberating enzymes in *S*. Tm and found that all *Salmonella enterica* genomes encode an uncharacterized gene with high sequence homology to characterized alpha – *N*- arabinofuranosidases. The Carbohydrate-Active Enzymes database (CAZy; http://www.cazy.org), predicts *S*. Tm to encode a single arabinofuranosidase of the 48 predicted glycoside hydrolases in *S*. Tm SL1344 (Table S1). A multiple sequence alignment program (MAFT) was used to align the *S*. Tm arabinofuranosidase sequence to functionally characterized arabinofuranosidases (Figure S5A) (Cartmell *et al*., 2011). The predicted structure of this uncharacterized hydrolase using Phyre revealed that the critical residues meditating substrate binding and enzymatic activity are conserved between this protein and previously characterized alpha – *N*- arabinofuranosidases (Figure 5A). Interestingly, our transcriptome data revealed that this putative hydrolase (STM0148) was differentially expressed *in vivo* (Figure 5B), strongly suggesting that *S*. Tm expresses this enzyme during gut colonization.

**Figure 5.**
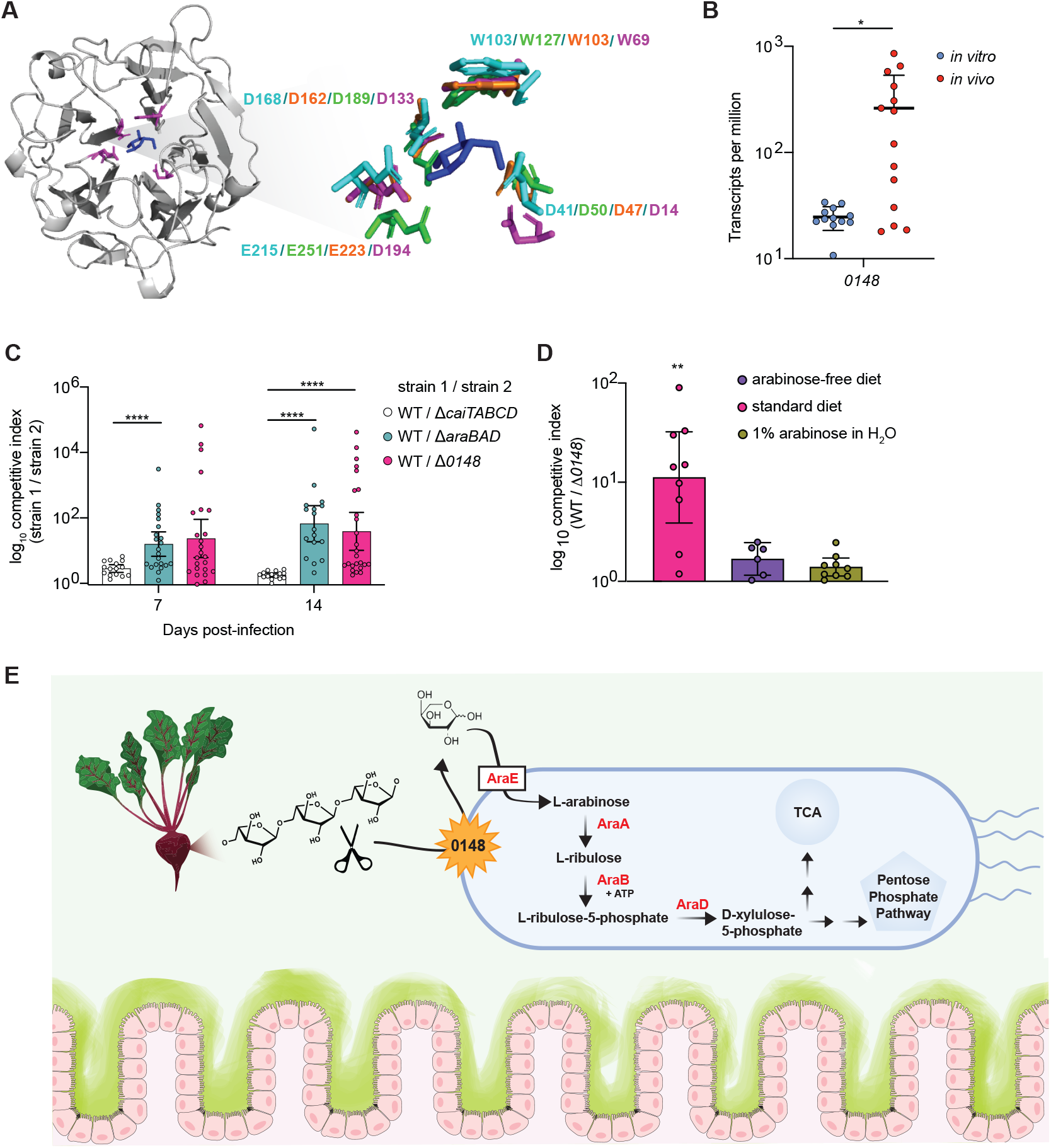
*S*. Tm uses a functionally conserved arabinofuranosidase to colonize the gut. A) Predicted structure of STMO148 generated with Phyre. Critical residues highlightedin purple surround L-arabinose. Call-out shows structural alignment of critical residues from *S*. Tm (in purple) with characterized arabinofuranosidase enzymes from *Cellvibrio japonicus* (PDB accession 3QEE) in teal, *Bacillus subtilis* in green (PDB accession 3C7G), and *Streptomyces avermitilis* (PDB accession 3AKH) in orange. L-arabinose molecule is derived from the *S. avermitilis* structure (3AKH). B) *Salmonella* Typhimurium putative arabinofuranosidase (STM0148) is differentially expressed *in vivo* compared to *in vitro*. T test *p < 0.05. C) 129X1/SvJ mice were inoculated with an equal mixture of the indicated strains. The competitive index from the feces was determined at 7- and 14-days post – infection. Each dot represents data from one animal (biological replicate). Bars represent +-SEM. *p < 0.05, ** p < 0.01, *** p < 0.001, **** p < 0.0001. Mann – Whitney U D) 129X1/SvJ mice were inoculated with an equal mixture of WT STm and 0148. Exogenous L-arabinose was provided in the drinking water of the indicated group shown in green, and an arabinose-free diet was given to the indicated group shown in purple. The competitive index was determined at 7 DPI. Each dot represents data from one animal (biological replicate). Bars represent +-SEM. *p < 0.05, ** p < 0.01. Kruskall-Wallis E) Graphical model of the proposed mechanism of L-arabinose acquisition and subsequent utilization. See also Figure S5 and Table S1.

To test if the function of this putative arabinofuranosidase contributes to colonization of the gut, we engineered a strain lacking this gene (*STM0148*). We then infected 129X1/SvJ mice with 1:1 mixtures of WT and *D0148::Kan^R^* or the *DcaiTABCD::Kan^R^* control *S*. Tm strains, and calculated the CI. In these experiments, mice were maintained on a standard chow which contains L-arabinose-containing plant polysaccharides in the form of crude plant material. WT *S. Tm* was recovered significantly more than *D0148 S*. Tm over a time course of infection (Figure 5C), indicating this putative arabinofuranosidase confers a competitive advantage for *S. Tm in vivo*.

To further interrogate the role of *S*. Tm’s putative arabinofuranosidase in the acquisition and utilization of L-arabinose *in vivo*, we infected 129X1/SvJ mice with a 1:1 ratio of WT and *D0148::Kan^R^ S*. Tm strains and fed mice a polysaccharide defined diet free of L-arabinose-containing sugars. WT and *D0148::Kan^R^ S*. Tm strains were recovered at similar levels in the L-arabinose-deficient guts, indicating that the arabinofuranosidase is not advantageous to *S*. Tm when mice are fed a diet lacking L-arabinose-containing polysaccharides (Figure 5D). To further test this notion, we infected a group of 129X1/SvJ mice fed a standard diet and given water supplemented with 1% L-arabinose with equal concentrations of the WT and *D0148::Kan^R^ S*. Tm strains. In these mice, the WT strain did not have a competitive advantage compared to *D0148::Kan^R^ S*. Tm, suggesting that both the removal of L-arabinose containing polysaccharides and the supplementation of free L-arabinose were sufficient to ablate the competitive advantage conferred by the arabinofuranosidase (Figure 5D). Together these data indicate that *S*. Tm utilizes a functional arabinofuranosidase *in vivo*, where the enzyme liberates free L-arabinose from complex plant polysaccharides that allow *S*. Tm to expand in the guts of superspreader mice (Figure 5E).

## DISCUSSION

Superspreaders are responsible for most disease spread of *Salmonella*, yet the molecular mechanisms that underlie superspreader development are understudied. The emergence of superspreaders likely depends on a number of complex interactions between host, microbiome, and pathogen that appear to be driven by the metabolic microenvironment of the intestine (Gopinath et al., 2012; Jacobson et al., 2018; Lawley et al., 2008). Here, we leveraged a mouse model of chronic *Salmonella* infection in which the gut ecosystem is not perturbed by antibiotics. Using this model, we identified mechanisms by which superspreaders arise in mice infected with *S*. Tm (Monack, Bouley and Falkow, 2004; Lawley *et al*., 2006; Gopinath, Carden and Monack, 2012). The use of unbiased fecal-metabolome mapping of mice harboring heterogenous levels of *S*. Tm over time allowed us to resolve a metabolic phenotype of *Salmonella* superspreader mice. In parallel, we performed transcriptomics on *S*. Tm recovered from the feces of superspreader mice. By analyzing these two datasets in tandem, we identified L-arabinose metabolism as a critical feature of superspreading mice, revealing that the consumption of this pentose sugar was advantageous for *S*. Tm’s expansion in the gut. Collectively, our data shed light on a nutrient-driven mechanism by which *Salmonella* expands within the guts of superspreading mice and highlights the role that diet-derived factors, such as L-arabinose, may play in superspreading hosts.

To colonize the gut, *Salmonella* must overcome colonization resistance while employing mechanisms of expansion in parallel. In addition to competing with *Salmonella* for carbohydrate sources, commensals can interfere with *Salmonella* expansion through production of short chain fatty acids or antimicrobial peptides (Jacobson *et al*., 2018; Sorbara and Pamer, 2019). The microbiota can also indirectly inhibit *Salmonella* through induction of innate and adaptive immune responses (Chung *et al*., 2012). Concurrently, *Salmonella* has a type VI secretion system it employs for contact-dependent killing of commensals (Sana *et al*., 2016). *Salmonella* also can exploit commensals by using their fermentation end-product, hydrogen, as an electron donor (Maier *et al*., 2013). Here we propose an additional mechanism of pathogen expansion that relies on the selective consumption of a dietary sugar, L-arabinose.

Rolf Freter’s nutrient niche theory claims that a pathogen can only colonize an ecological niche if it is more efficient at using a particular limiting nutrient than its competitors (Freter et al.,1983). It is well established that the ability to degrade carbohydrates, diet or host-mucus derived, provides a competitive advantage to colonizing microbes of the gut (Martens et al., 2014; Wardman et al., 2022). The carbohydrate active enzymes (CAZymes) involved in the degradation of complex carbohydrates include glycoside hydrolases, polysaccharide lyases, and carbohydrate esterases. CAZymes tend to make up large proportions of gut commensal genomes. For example, the common gut resident Bacteroides *thetaiotaomicron* dedicates ~7% of its genome to glycoside hydrolases (Kaoutari *et al*., 2013). While CAZymes have been studied in detail within various commensal microbes and marine ecosystems (Sichert *et al*., 2020; Solanki *et al*., 2022), the role these enzymes play in pathogenic bacteria are still unclear. Interestingly, chitinases, which hydrolyze chitin polymers into N-acetyl glucosamine oligomers, are a well-studied family of glycoside hydrolases in pathogens. Recent work has characterized *Salmonella’s* GH18 family chitinases and have shown them to be important for epithelial adhesion and invasion (Devlin *et al*., 2021; Chandra *et al*., 2022). Chitinases have been identified as important virulence factors for several other pathogenic bacterial species including *Legionella pneumophila* (Rehman *et al*., 2020), *Listeria monocytogenes* (Chaudhuri *et al*., 2013), *Vibrio cholerae* (Mondal *et al*., 2014), and Adherent-invasive *Escherichia coli* (Low *et al*., 2013). In contrast to chitinases mediating the process of infection as a virulence factor, we propose a mechanism in which *Salmonella* gains a selective advantage through the acquisition of dietary L-arabinose using a previously uncharacterized alpha-n-arabinofuranosidase. We show *Salmonella’s* alpha-n-arabinofuranosidase shares critical residues with functionally characterized arabinofuranosidase enzymes in the glycoside hydrolase family 43 (Figure 5A-5B) (Cartmell *et al*., 2011). We also provide evidence that *Salmonella’s* alpha-n-arabinofuranosidase is functionally active *in vivo* and confers a competitive advantage for *Salmonella* in the gut (Figure 5).

Metabolic cross-feeding amongst members of the microbiota can result in complex food chains, where species may depend on others for partial glycan degradation (Koropatkin *et al*., 2012). For example, *Bacteroides* species can act as primary degraders of complex carbohydrates, releasing oligosaccharides for secondary degraders to process and consume (Fischbach and Sonnenburg, 2011). Cross-feeding examples have been described for various dietary glycans including arabinoxylan (Rogowski *et al*., 2015) and inulin (Rakoff-Nahoum *et al*., 2016). *S*. Tm is known to depend on commensal bacteria for liberation of readily consumable monosaccharides like fucose (Ng *et al*., 2013). We speculate that as *S*. Tm becomes a more dominant member of the community, self-reliance is a failsafe strategy. *S*. Tm using its arabinofuranosidase to acquire L-arabinose may be an example of such an autonomous pathway. This study provides the groundwork for future research to address the role of *Salmonella’s* arabinofuranosidase in the metabolic cross-feeding webs occurring in the gut.

Overall, we have leveraged both metabolomics and transcriptomics of *Salmonella*-infected superspreader mice to define an important molecular mechanism of pathogen expansion within the gastrointestinal tract. These results shed light on the metabolomic landscape within superspreaders and provide a holistic snapshot of complex host-pathogen interactions occurring in the gut, at the resolution of single metabolites. The metabolomic and transcriptomic datasets presented in this study will likely enable future studies of additional mechanisms that underlie the formation of superspreaders. Ultimately this work contributes to the understanding of pathogen expansion within superspreader hosts, an area of study necessary for disease transmission control.

## Supporting information

Key resources table

metabolomics data

## Acknowledgements

The authors thank Manuel Amieva, and all members of the Monack and Amieva laboratories for valuable discussions and review of the manuscript. Authors also thank Justin Sonnenburg, Steven Higginbottom, and all personnel in the gnotobiotic group at Stanford University for their expertise and generosity. Research reported in this publication was supported by grants R01-AI116059 and R01AI095396 from the National Institute of Allergy and Infectious Diseases, United States (D.M.M.), Paul Allen Stanford Discovery Center on Systems Modeling of Infection (to D.M.M.), Gates Grand Challenge Grant from Bill & Melinda Gates Foundation (to D.M.M.), and the Graduate Research Fellowship Program funded by the National Science Foundation (S.R.). The content is solely the responsibility of the authors and does not necessarily represent the official views of the National Institutes of Health.

## Author Contributions

Conceptualization, S.R. and D.M.M.; Investigation, S.R., L.M., and A.C.; Writing — Original Draft, S.R. and D.M.M.; Writing — Review & Editing, all authors; Funding Acquisition, S.R. and D.M.M.

## STAR Methods

### Key Resources Table

Separate file

### Resource Availability

#### Lead Contact and Materials Availability

Further information and requests for resources and reagents should be directed to the Lead Contact, Denise M. Monack.

#### Data and code availability

- Untargeted metabolomics data are attached as a supplementary file to this manuscript.
- Raw RNA-seq data discussed in this study have been deposited in NCBI’s Gene Expression Omnibus (Edgar *et al*., 2002) and are accessible through GEO Series accession number GSE207764 (https://www.ncbi.nlm.nih.gov/geo/query/acc.cgi?acc=GSE207764).
- Code is available at https://github.com/sruddle/superspreader

### Experimental Model and Subject Details

#### Ethics Statement

Experiments involving animals were performed in accordance with NIH guidelines, the Animal Welfare Act, and US federal law. All animal experiments were approved by the Stanford University Administrative Panel on Laboratory Animal Care (APLAC) and overseen by the Institutional Animal Care and Use Committee (IACUC) under Protocol ID 12826. Animals were housed in a centralized research animal facility accredited by the Association of Assessment and Accreditation of Laboratory Animal Care (AAALAC) International.

#### Mouse Strains and Husbandry

129X1/SvJ mice were obtained from Jackson Laboratories. Female mice (6-7 weeks old) were housed under specific pathogen-free conditions in filter-top cages that were changed bi-monthly by veterinary or research personnel. Sterile water and food were provided ad libitum. Mice were given at least one week to acclimate to the Stanford Animal Biohazard Research Facility prior to experimentation. Mouse age, sex, lineage, and source facility were tracked for all experiments and did not correlate with infection outcomes.

Swiss-Webster mice were used for gnotobiotic experiments, and the sterility of germ-free mice was verified through routine 16S PCR amplification and anaerobic culture of feces by gnotobiotic facility staff. Mice were maintained on a 12-h light/dark cycle at 20.5 °C at ambient humidity, fed ad libitum, and maintained in flexible film gnotobiotic isolators for the duration of all experiments. Mice were controlled for sex and age within experiments.

#### Bacterial Strains and Growth Conditions

S. Typhimurium SL1344 strains were maintained aerobically on LB agar supplemented with 200 g/mL streptomycin (LB-strep), 200 g/mL streptomycin + 15 g/mL tetracycline (LB-strep-tet), or 200 g/mL streptomycin + 40 g/mL kanamycin (LB-strep-kan) and grown aerobically overnight at 37°C with aeration.

### Method Details

#### Mouse Infections

Mice were infected with 10^8^ CFU S. Typhimurium by drinking. For competitive index experiments, mice were infected with a 1:1 mixture of each S. Typhimurium strain by drinking, totaling 10^8^ CFU S. Typhimurium. 1-2 fecal pellets were collected and resuspended in 500μl PBS. To determine CFU/g feces, fecal pellets were weighed, serially diluted, and plated onto LB plates supplemented with appropriate antibiotics. Competitive Index (CI) values were calculated as previously described (Beuzón and Holden, 2001).

For terminal experiments, mice were sacrificed at the indicated time points. Mice were euthanized with carbon dioxide. Sterile dissection tools were utilized to collect spleen, liver, mesenteric lymph nodes (MLN), small intestine, cecum, and colon. Tissues were collected in 1-3 mL PBS and homogenized. Homogenates were plated as serial dilutions on LB agar supplemented with 200 mg/mL of appropriate antibiotics to enumerate CFU/g tissue. Each experiment was conducted a minimum of 3 times to ensure reproducibility.

### Untargeted Metabolomics

#### Sample collection

Untargeted Metabolomic profiling on fecal samples was conducted using ultra-high-performance liquid chromatography-tandem mass-spectrometry by Metabolon Inc. (Morrisville, USA). In summary, fecal samples (2-3 fecal pellets) were collected before infection and on days 7,14, and 21 after infection (with 10^8^ CFU S. Typhimurium). Mice were single housed for the fecal pellet collection. Samples were immediately transferred to dry ice and subsequently stored at −80°C until sent to Metabolon, Inc for processing and analysis.

#### Ultrahigh Performance Liquid Chromatography-Tandem Mass Spectroscopy (UPLC-MS/MS)

All methods utilized a Waters ACQUITY ultra-performance liquid chromatography (UPLC) and a Thermo Scientific Q-Exactive high resolution/accurate mass spectrometer interfaced with a heated electrospray ionization (HESI-II) source and Orbitrap mass analyzer operated at 35,000 mass resolution. The sample extract was dried then reconstituted in solvents compatible to each of the four methods. Each reconstitution solvent contained a series of standards at fixed concentrations to ensure injection and chromatographic consistency. One aliquot was analyzed using acidic positive ion conditions, chromatographically optimized for more hydrophilic compounds. In this method, the extract was gradient eluted from a C18 column (Waters UPLC BEH C18-2.1×100 mm, 1.7 μm) using water and methanol, containing 0.05% perfluoropentanoic acid (PFPA) and 0.1% formic acid (FA). Another aliquot was also analyzed using acidic positive ion conditions; however, it was chromatographically optimized for more hydrophobic compounds. In this method, the extract was gradient eluted from the same afore mentioned C18 column using methanol, acetonitrile, water, 0.05% PFPA and 0.01% FA and was operated at an overall higher organic content. Another aliquot was analyzed using basic negative ion optimized conditions using a separate dedicated C18 column. The basic extracts were gradient eluted from the column using methanol and water, however with 6.5mM Ammonium Bicarbonate at pH 8. The fourth aliquot was analyzed via negative ionization following elution from a HILIC column (Waters UPLC BEH Amide 2.1×150 mm, 1.7 μm) using a gradient consisting of water and acetonitrile with 10mM Ammonium Formate, pH 10.8.

#### Data Extraction and Compound Identification

Raw data was extracted, peak-identified and QC processed using Metabolon’s hardware and software. Compounds were identified by comparison to library entries of purified standards or recurrent unknown entities. Metabolon maintains a library based on authenticated standards that contains the retention time/index (RI), mass to charge ratio (*m/z*), and chromatographic data (including MS/MS spectral data) on all molecules present in the library. Furthermore, biochemical identifications are based on three criteria: retention index within a narrow RI window of the proposed identification, accurate mass match to the library +/- 10 ppm, and the MS/MS forward and reverse scores between the experimental data and authentic standards. The MS/MS scores are based on a comparison of the ions present in the experimental spectrum to the ions present in the library spectrum. While there may be similarities between these molecules based on one of these factors, the use of all three data points can be utilized to distinguish and differentiate biochemicals. More than 3300 commercially available purified standard compounds have been acquired and registered into LIMS for analysis on all platforms for determination of their analytical characteristics. Additional mass spectral entries have been created for structurally unnamed biochemicals, which have been identified by virtue of their recurrent nature (both chromatographic and mass spectral).

#### Statistical analysis of metabolomic data

Metabolon, USA performed normalization to sample volume, log transformation, and imputation of missing values, if any, with the minimum observed value for each compound. Analysis by two-way ANOVA with repeated measures identified biochemicals exhibiting significant interaction and main effects for experimental parameters of shedding level, time point, and infection status. Standard statistical analyses were performed in ArrayStudio on log transformed data by Metabolon, USA. PCA was generated using log transformed metabolite levels. Pathway analysis was performed using the XNomial package v1.0.4 in R v4.0.2 for likelihood ratio test with Benjamini & Hochberg correction.

#### RNA Isolation

RNA from cecal contents and feces was extracted using the QIAGEN RNeasy PowerMicrobiome Kit following manufacturer’s instructions. RNA from liquid cultures was extracted with the QIAGEN RNeasy Mini Kit following the QIAGEN supplementary protocol: purification of total RNA from bacteria. Three milliliters of culture were centrifuged at 6,000g for 10 min, resuspended in kit buffer RLT + beta-mercaptoethanol, and bead-beat using acid-washed glass beads for 5 min at 4°C. RNA was quantified using a Quibit and quality determined by an Agilent bioanalyzer.

#### RNA Library preparation and sequencing

RNA libraries were prepared using the Illumina stranded total RNA prep ligation with ribo-zero plus kit. rRNA was depleted prior to library construction and confirmed with Agilent Bioanalyzer Prokaryote Total RNA Pico. RNA integrity was confirmed with Agilent Bioanalyzer Prokaryote Total RNA Pico. *In vivo* and *in vitro* libraries underwent quality control by Novogene Corporation Inc. Pulled *in vivo and in vitro* libraries were loaded onto separate lanes of an S4 flow cell and sequenced on an Illumina NovaSeq 6000 using the PE150 strategy. Data were unequally shared, 10.11G raw data (100.4 million paired reads) for each *in vivo* sample and 4.5G raw data (15 million paired reads) for each *in vitro* sample.

#### RNA-seq analysis

RNA-seq analysis was performed using the nf-core RNA-seq pipeline development branch version (Ewels *et al*., 2020). Quality control was performed using FastQC v0.11.9. Adapters and low-quality reads were filtered by TrimGalore! with default parameters v0.6.5. Quality reads were aligned to *S*. Tm strain SL1344 genome using HiSAT2 v2.2.1 (Kim *et al*., 2019). Stringtie was used to assemble transcript alignments and for quantification v2.1.4 (Pertea *et al*., 2015). Reads were further processed in Deseq2 (Love *et al*., 2014) for differential expression analysis. Results were plotted in Graph pad prism v8.1.2 or R using ggplot2. Biocyc Pathway Tools was used for pathway enrichment analysis (Karp *et al*., 2019).

#### Bacterial Strain Construction

Genes in the SL1344 genome were targeted for deletion following lambda red mutagenesis protocol (Datsenko and Wanner, 2000). Briefly, primers were used to amplify the kanamycin resistance cassette of pKD4, and PCR products were purified. *S*. Tm strains 14028 containing pKD46 were grown at 30°C, 200 rpm to mid-log phase, L-arabinose was added to a final concentration of 50 mM and bacteria were incubated for an additional hour. 1-5 μg of purified PCR product was added to washed and concentrated bacteria. Bacteria were electroporated and recovered in 30 °C SOC broth for 3 hours with shaking. Cultures were plated on antibiotic selection plates and grown overnight at 42 °C to cure cells of the pKD46 plasmid. Colonies were PCR verified for the kanamycin resistance insertion and endogenous gene disruption. Removal of the pKD46 plasmids was confirmed by a loss of carbenicillin resistance. P-22 phage transduction was used to introduce desired gene knockouts to SL1344. H-5 phage was used to confirm final stains as lysogen-free. Final Sl1344 strains were confirmed through PCR.

#### Custom research diets

Animals were maintained on standard diet (SD, Envigo TD.2018, Teklad Global 2018 rodent diet) for at least one week prior to experimentation. If mice were switched to an arabinose-free diet (polysaccharide-defined: custom diet, Envigo TD.150689 all carbohydrate and fiber sources replaced with dextrose monohydrate), this occurred at time of infection.

#### L-Arabinose Quantification

Feces were weighed, homogenized in 10% ethanol, and spun at 14,000g for 5 minutes. Supernatant was sent to UCSD’s Glycoanalytics Core. There, Samples were centrifuged at 10,000g for 5 minutes and spin filtered using prewashed 3K spin filtration unit (Centrifugal device, Pall, Life Sciences, Part No. OD003C34). Flow through was spiked with 1.0μg of Inositol as internal standard and lyophilized. Dried material was reduced using sodium borohydride in presence of 1M ammonium hydroxide, overnight at room temperature. Excess reducing agent was neutralized on ice bath using 30% aqueous acetic acid solution. Samples were then dried by nitrogen flush and co-evaporated with 9:1 methanol: acetic acid mixture (3 times) followed by anhydrous methanol (3 times). Finally, the samples were acetylated using mixture of pyridine and acetic anhydride (1:1 v/v) and alditol acetylate of monosaccharides were analyzed by GCMS (Agilent Technologies, 7820 GC System attached with 5975-MSD). Restek-5ms column (30m x 0.25mm x 0.25μm) was used for profiling of the monosaccharides, Ultrapure He was used as carrier gas in split less mode at a flow rate of 1.2mL/min. Quantification of monosaccharides were done based on the response factors obtained from known amount of standard mixture.

#### Lipocalin ELISA

Feces were collected from mice, weighed, and stored at −80 °C until protein quantification could be performed. Samples were homogenized by bead beating and protein was assessed using a Lipocalin-2 (LCN2) Mouse ELISA Kit (Invitrogen) according to manufacturer’s instructions.

#### qRT-PCR

Spleen and colon tissues were harvested from mice and stored at −80 °C until RNA extractions were performed. Tissues were homogenized by bead beating and RNA was isolated using an RNeasy Mini Kit (Qiagen). RNA concentration was determined with a Nanodrop spectrophotometer and samples were stored at −80 °C. cDNA was synthesized from RNA samples using a SuperScript III First-Strand Synthesis Kit (Thermo Fisher Scientific) and stored at −20 °C. To assess expression of inflammatory genes, cDNA and gene-specific primers were used in combination with FastStart Universal SYBR Green Master Mix (Sigma Aldrich) to amplify transcripts.

#### Quantification and Statistical Analysis

All statistical analyses were performed in R or Prism v. 8.1.2 (GraphPad) and visualized with ggplot2 and Prism v. 8.1.2 (GraphPad). Statistical significance of all CFU counts were determined by 2-way ANOVA in Prism v. 8.1.2 (GraphPad). Statistical significance of time-course competitive index infection data was determined by Mann-Whitney U test in Prism v. 8.1.2 (GraphPad).

---

**Supplemental figure 1. *S*. Tm superspreaders and non-superspreaders have differentially enriched metabolic pathways at 7 and 14 DPI**

Corresponds with figure 1

A) Differential pathway enrichment of non-superspreaders and superspreaders at 7 DPI with multiple hypothesis correction (Likelihood ratio with Benjamini & Hochberg correction).

B) Differential pathway enrichment of non-superspreaders and superspreaders at 21 DPI with multiple hypothesis correction (Likelihood ratio with Benjamini & Hochberg correction).

**Supplemental figure 2. *S*. Tm genetic pathways enriched *in vivo* and *in vitro***

Corresponds with figure 2

A) Log2 fold differences of differentially enriched genetic pathways (adjusted p-value < 0.05) *in vitro* (blue) and *in vivo* (red). Each data point represents single gene within each sub pathway (y-axis). Sub pathways are annotated with corresponding major pathways (brackets along y-axis)

B) Principal component analysis of *in vivo* and *in vitro* transcriptomes. *In vivo* and *in vitro* transcriptomes separate along PC1.

**Supplemental figure 3: Exogenous dietary L-arabinose does not induce an inflammation**

Corresponds with figure 4

A) Lipocalin-2 levels detected in feces of the indicated groups 4-days post treatment (diet and/or infection) by Elisa. N=5 mice

B) qRT-PCR of mRNA extracted from colon tissue from indicated groups 5 days post treatment. N=5 mice

C) qRT-PCR of mRNA extracted from spleens from indicated groups 5 days post treatment. N=5 mice

**Supplemental figure 4: Carnitine supplementation does not confer a competitive advantage to WT *S*. Tm and L-arabinose concentrations decrease during infection in germ-free mice**

Corresponds with figure 4

A) 129X1/SvJ mice were inoculated with an equal mixture of WT *S*. Tm and *DcaiTABCD*. Each dot represents data from one animal (biological replicate). Bars represent +-SEM. *p < 0.05, ** p < 0.01. Kruskal-Wallis. No difference across groups. N=10 per group.

B) L-Arabinose concentration in the germ-free mouse feces was quantified by liquid LC/MS. Students T test, *** p < 0.001

**Supplemental figure 5. Salmonella’s arabinofuranosidase has conserved functional residues with previously characterized arabinofuranosidases**

Corresponds with figure 5

A) Amino acid sequence alignment of STM0148 (grey highlight) against three characterized arabinofuranosidase amino acid sequences. Red rectangles show conserved critical residues for enzyme function.

## REFERENCES

Ali, M. M. et al. (2014) ‘Fructose-Asparagine Is a Primary Nutrient during Growth of Salmonella in the Inflamed Intestine’, PLoS Pathog, 10(6), p. 1004209. doi: 10.1371/journal.ppat.1004209.

Ammar, E. M., Wang, X. and Rao, C. V (no date) ‘Regulation of metabolism in Escherichia coli during growth on mixtures of the non-glucose sugars: arabinose, lactose, and xylose OPEN’. doi: 10.1038/s41598-017-18704-0.

Barrett, E. L. and Riggs, D. L. (1982) ‘Evidence of a second nitrate reductase activity that is distinct from the respiratory enzyme in Salmonella typhimurium’, Journal of bacteriology. J Bacteriol, 150(2), pp. 563–571. doi: 10.1128/JB.150.2.563-571.1982.

Barthel, M. et al. (2003) ‘Pretreatment of mice with streptomycin provides a Salmonella enterica serovar Typhimurium colitis model that allows analysis of both pathogen and host’, Infect. Immun. American Society for Microbiology, 71(5), pp. 2839–2858. doi: 10.1128/iai.71.5.2839-2858.2003.

Beuzón, C. R. and Holden, D. W. (2001) ‘Use of mixed infections with Salmonella strains to study virulence genes and their interactions in vivo’, Microbes and infection. Microbes Infect, 3(14-15), pp. 1345–1352. doi: 10.1016/S1286-4579(01)01496-4.

Cartmell, A. et al. (2011) ‘The structure and function of an arabinan-specific α-1,2-arabinofuranosidase identified from screening the activities of bacterial GH43 glycoside hydrolases’, Journal of Biological Chemistry, 286(17), pp. 15483–15495. doi: 10.1074/jbc.M110.215962.

Chakravorty, M. (1964) ‘Induction and repression of L-arabinose isomerase in salmonella typhimurium’, BBA - Enzymological Subjects, 85(1), pp. 152–161. doi: 10.1016/0926-6569(64)90175-0.

Chandra, K. et al. (2022) ‘GH18 family glycoside hydrolase Chitinase A of Salmonella enhances virulence by facilitating invasion and modulating host immune responses.’, PLoS pathogens, 18(4), p. e1010407. doi: 10.1371/journal.ppat.1010407.

Chaudhuri, S. et al. (2013) ‘The Listeria monocytogenes ChiA chitinase enhances virulence through suppression of host innate immunity’, mBio, 4(2). doi: 10.1128/mBio.00617-12.

Chessa, D. et al. (2008) ‘Salmonella enterica serotype Typhimurium Std fimbriae bind terminal a(1,2)fucose residues in the cecal mucosa’. doi: 10.1111/j.1365-2958.2008.06566.x.

Chung, H. et al. (2012) ‘Gut Immune Maturation Depends on Colonization with a Host-Specific Microbiota’, Cell. Cell Press, 149(7), pp. 1578–1593. doi: 10.1016/J.CELL.2012.04.037.

Collins, F. M. and Carter, P. B. (1978) ‘Growth of salmonellae in orally infected germfree mice’, Infection and Immunity, 21(1), pp. 41–47. doi: 10.1128/iai.21.1.41-47.1978.

Crum Cianflone, N. F. (no date) ‘Salmonellosis and the GI Tract: More than Just Peanut Butter’.

Datsenko, K. A. and Wanner, B. L. (2000) ‘One-step inactivation of chromosomal genes in Escherichia coli K-12 using PCR products’, Proceedings of the National Academy of Sciences of the United States of America, 97(12), pp. 6640–6645. doi: 10.1073/PNAS.120163297.

Deatherage Kaiser, B. L. et al. (2013) ‘A Multi-Omic View of Host-Pathogen-Commensal Interplay in Salmonella-Mediated Intestinal Infection’, PLOS ONE. Public Library of Science, 8(6), p. e67155. doi: 10.1371/JOURNAL.PONE.0067155.

Devlin, J. R. et al. (2021) ‘Salmonella enterica serovar Typhimurium chitinases modulate the intestinal glycome 1 and promote small intestinal invasion’, bioRxiv, p. 2021.12.06.471358. doi: 10.1371/journal.ppat.1010167.

Diard, M. et al. (2013) ‘Stabilization of cooperative virulence by the expression of an avirulent phenotype’, Nature 2013 494:7437. Nature Publishing Group, 494(7437), pp. 353–356. doi: 10.1038/nature11913.

Faber, F. et al. (2017) ‘Respiration of Microbiota-Derived 1,2-propanediol Drives Salmonella Expansion during Colitis’, PLOS Pathogens. Public Library of Science, 13(1), p. e1006129. doi: 10.1371/JOURNAL.PPAT.1006129.

Fischbach, M. A. and Sonnenburg, J. L. (2011) ‘Eating For Two: How Metabolism Establishes Interspecies Interactions in the Gut’, Cell Host & Microbe. Cell Press, 10(4), pp. 336–347. doi: 10.1016/J.CHOM.2011.10.002.

Freter, R. et al. (1983) ‘Mechanisms that control bacterial populations in continuous-flow culture models of mouse large intestinal flora’, Infection and Immunity, 39(2), pp. 676–685. doi: 10.1128/iai.39.2.676-685.1983.

Gopinath, S. et al. (no date) ‘Role of disease-associated tolerance in infectious superspreaders’. doi: 10.1073/pnas.1409968111.

Gopinath, S., Carden, S. and Monack, D. (2012a) ‘Shedding light on Salmonella carriers’, Trends in Microbiology, 20, pp. 320–327. doi: 10.1016/j.tim.2012.04.004.

Gopinath, S., Carden, S. and Monack, D. (2012b) ‘Shedding light on Salmonella carriers’, Trends in Microbiology. doi: 10.1016/j.tim.2012.04.004.

Hockenberry, A. M. et al. (2021) ‘Microbiota-derived metabolites inhibit Salmonella virulent subpopulation development by acting on single-cell behaviors’, Proceedings of the National Academy of Sciences of the United States of America. National Academy of Sciences, 118(31). doi: 10.1073/PNAS.2103027118/SUPPL_FILE/PNAS.2103027118.SM02.MOV.

Jacobson, A. et al. (2018a) ‘A Gut Commensal-Produced Metabolite Mediates Colonization Resistance to Salmonella Infection’, Cell Host and Microbe, 24(2). doi: 10.1016/j.chom.2018.07.002.

Jacobson, A. et al. (2018b) ‘A Gut Commensal-Produced Metabolite Mediates Colonization Resistance to Salmonella Infection’, Cell Host and Microbe. Cell Press, 24(2), pp. 296–307.e7. doi: 10.1016/j.chom.2018.07.002.

Kaoutari, A. El et al. (2013) ‘The abundance and variety of carbohydrate-active enzymes in the human gut microbiota’, Nature Reviews Microbiology. Nature Publishing Group, 11(7), pp. 497–504. doi: 10.1038/nrmicro3050.

Karp, P. D. et al. (2019) ‘The BioCyc collection of microbial genomes and metabolic pathways’, Briefings in bioinformatics. Brief Bioinform, 20(4), pp. 1085–1093. doi: 10.1093/BIB/BBX085.

Kim, D. et al. (2019) ‘Graph-based genome alignment and genotyping with HISAT2 and HISAT-genotype’, Nature Biotechnology 2019 37:8. Nature Publishing Group, 37(8), pp. 907–915. doi: 10.1038/s41587-019-0201-4.

Koropatkin, N. M., Cameron, E. A. and Martens, E. C. (2012) ‘How glycan metabolism shapes the human gut microbiota’, Nature Reviews Microbiology. Nature Publishing Group, 10(5), pp. 323–335. doi: 10.1038/nrmicro2746.

Lawley, T. D. et al. (2006) ‘Genome-Wide Screen for Salmonella Genes Required for Long-Term Systemic Infection of the Mouse’, 2(2). doi: 10.1371/journal.ppat.0020011.

Lawley, Trevor D. et al. (2008) ‘Host transmission of Salmonella enterica serovar Typhimurium is controlled by virulence factors and indigenous intestinal microbiota’, Infect. Immun., 76(1), pp. 403–416. doi: 10.1128/iai.01189-07.

Lawley, Trevor D et al. (2008) ‘Host Transmission of Salmonella enterica Serovar Typhimurium Is Controlled by Virulence Factors and Indigenous Intestinal Microbiota’, INFECTION AND IMMUNITY, 76(1), pp. 403–416. doi: 10.1128/IAI.01189-07.

López-Garrido, J. et al. (2015) ‘Virulence gene regulation by L-arabinose in Salmonella enterica’, Genetics. Genetics Society of America, 200(3), pp. 807–819. doi: 10.1534/GENETICS.115.178103/-/DC1.

Lopez, C. A. et al. (2012) ‘Phage-mediated acquisition of a type III secreted effector protein boosts growth of salmonella by nitrate respiration’, mBio. mBio, 3(3). doi: 10.1128/MBIO.00143-12.

Love, M. I., Huber, W. and Anders, S. (2014) ‘Moderated estimation of fold change and dispersion for RNA-seq data with DESeq2’, Genome Biology, 15(12), pp. 1–21. doi: 10.1186/s13059-014-0550-8.

Low, D. et al. (2013) ‘Chitin-binding domains of Escherichia coli ChiA mediate interactions with intestinal epithelial cells in mice with colitis’, Gastroenterology. Gastroenterology, 145(3). doi: 10.1053/J.GASTRO.2013.05.017.

Maier, L. et al. (2013) ‘Microbiota-derived hydrogen fuels salmonella typhimurium invasion of the gut ecosystem’, Cell Host and Microbe, 14(6), pp. 641–651. doi: 10.1016/j.chom.2013.11.002.

Martens, E. C. et al. (2009) ‘Complex glycan catabolism by the human gut microbiota: The bacteroidetes sus-like paradigm’, Journal of Biological Chemistry. Elsevier, 284(37), pp. 24673–24677. doi: 10.1074/JBC.R109.022848/ATTACHMENT/34E0EC60-0F4F-46C3-9EB0-BD301F854F0B/MMC1.PDF.

Martens, E. C. et al. (2011) ‘Recognition and Degradation of Plant Cell Wall Polysaccharides by Two Human Gut Symbionts’, PLOS Biology. Public Library of Science, 9(12), p. e1001221. doi: 10.1371/JOURNAL.PBIO.1001221.

Martens, E. C. et al. (2014a) ‘The devil lies in the details: How variations in polysaccharide fine-structure impact the physiology and evolution of gut microbes’, Journal of Molecular Biology. Academic Press, pp. 3851–3865. doi: 10.1016/j.jmb.2014.06.022.

Martens, E. C. et al. (2014b) ‘The devil lies in the details: How variations in polysaccharide fine-structure impact the physiology and evolution of gut microbes’, Journal of Molecular Biology. Elsevier Ltd, 426(23), pp. 3851–3865. doi: 10.1016/j.jmb.2014.06.022.

Mayer, C. and Boos, W. (2005) ‘Hexose/Pentose and Hexitol/Pentitol Metabolism’, EcoSal Plus. American Society for Microbiology, 1(2). doi: 10.1128/ECOSALPLUS.3.4.1/FORMAT/EPUB.

McGovern, C. M. and Leavitt, J. W. (1997) ‘Typhoid Mary: Captive to the Public’s Health.’, The Journal of American History, 84(1), p. 271. doi: 10.2307/2952837.

McLaughlin, P. A. et al. (2019) ‘Inflammatory monocytes provide a niche for Salmonella expansion in the lumen of the inflamed intestine’, PLOS Pathogens. Public Library of Science, 15(7), p. e1007847. doi: 10.1371/JOURNAL.PPAT.1007847.

Meadows, J. A. and Wargo, M. J. (2015) ‘Carnitine in bacterial physiology and metabolism’, Microbiology (United Kingdom), 161(6), pp. 1161–1174. doi: 10.1099/mic.0.000080.

Monack, D. M., Bouley, D. M. and Falkow, S. (2004) ‘Salmonella typhimurium Persists within Macrophages in the Mesenteric Lymph Nodes of Chronically Infected Nramp1 / Mice and Can Be Reactivated by IFN Neutralization’, The Journal of Experimental Medicine J. Exp. Med. □ The. Rockefeller University Press, 199(2), pp. 231–241. doi: 10.1084/jem.20031319.

Mondal, M. et al. (2014) ‘The Vibrio cholerae Extracellular Chitinase ChiA2 Is Important for Survival and Pathogenesis in the Host Intestine’, 9(9). doi: 10.1371/journal.pone.0103119.

Ng, K. M. et al. (2013a) ‘Microbiota-liberated host sugars facilitate post-antibiotic expansion of enteric pathogens’, Nature, 502(7469), pp. 96–99. doi: 10.1038/NATURE12503.

Ng, K. M. et al. (2013b) ‘Microbiota-liberated host sugars facilitate post-antibiotic expansion of enteric pathogens’, Nature. Nature Publishing Group, 502(7469), pp. 96–99. doi: 10.1038/nature12503.

Parry, C. M. et al. (2002) ‘Typhoid Fever’, New England Journal of Medicine, 347(22), pp. 1770–1782. doi: 10.1056/NEJMRA020201.

Pertea, M. et al. (2015) ‘StringTie enables improved reconstruction of a transcriptome from RNA-seq reads’, Nature Biotechnology 2015 33:3. Nature Publishing Group, 33(3), pp. 290–295. doi: 10.1038/nbt.3122.

Philip A. Ewels, Alexander Peltzer, Sven Fillinger, Harshil Patel, Johannes Alneberg, Andreas Wilm, Maxime Ulysse Garcia, P. D. T. & S. N. (2020) ‘The nf-core framework for community-curated bioinformatics pipelines’, Nature Biotechnology, 38(3), p. 271. doi: 10.1038/s41587-020-0435-1.

Rakoff-Nahoum, S., Foster, K. R. and Comstock, L. E. (2016) ‘The evolution of cooperation within the gut microbiota’, Nature 2015 533:7602. Nature Publishing Group, 533(7602), pp. 255–259. doi: 10.1038/nature17626.

Rehman, S. et al. (2020) ‘Structure and functional analysis of the Legionella pneumophila chitinase ChiA reveals a novel mechanism of metal-dependent mucin degradation’, PLOS Pathogens. Public Library of Science, 16(5), p. e1008342. doi: 10.1371/JOURNAL.PPAT.1008342.

Rivera-Chávez, F. and Bäumler, A. J. (2015) ‘The Pyromaniac Inside You: Salmonella Metabolism in the Host Gut’, Annual Review of Microbiology, 69(1), pp. 31–48. doi: 10.1146/annurev-micro-091014-104108.

Rogers, A. W. L., Tsolis, R. M. and Bäumler, A. J. (2021) ‘Salmonella versus the Microbiome’, Microbiology and Molecular Biology Reviews, 85(1), pp. 1–30. doi: 10.1128/mmbr.00027-19.

Rogowski, A. et al. (2015) ‘ARTICLE Glycan complexity dictates microbial resource allocation in the large intestine’, Nature Communications. doi: 10.1038/ncomms8481.

Sana, T. G. et al. (2016) ‘Salmonella Typhimurium utilizes a T6SS-mediated antibacterial weapon to establish in the host gut’, Proceedings of the National Academy of Sciences of the United States of America. National Academy of Sciences, 113(34), pp. E5044–E5051. doi: 10.1073/PNAS.1608858113/SUPPL_FILE/PNAS.201608858SI.PDF.

Schleif, R. (2021) ‘A Career’s Work, the L-Arabinose Operon: How It Functions and How We Learned It’. doi: 10.1128/ecosalplus.

Shelton, C. D. et al. (2022) ‘Salmonella enterica serovar Typhimurium uses anaerobic respiration to overcome propionate-mediated colonization resistance’, Cell Reports. Elsevier B.V., 38(1). doi: 10.1016/J.CELREP.2021.110180.

Sichert, A. et al. (2020) ‘Verrucomicrobia use hundreds of enzymes to digest the algal polysaccharide fucoidan’, Nature Microbiology. Springer US, 5(8), pp. 1026–1039. doi: 10.1038/s41564-020-0720-2.

Solanki, V. et al. (2022) ‘Glycoside hydrolase from the GH76 family indicates that marine Salegentibacter sp. Hel_I_6 consumes alpha-mannan from fungi’, ISME Journal. Springer US, (February). doi: 10.1038/s41396-022-01223-w.

Sommer, F. et al. (2017) ‘The resilience of the intestinal microbiota influences health and disease’. doi: 10.1038/nrmicro.2017.58.

Sonnenburg, E. D. and Sonnenburg, J. L. (2014) ‘Starving our microbial self: The deleterious consequences of a diet deficient in microbiota-accessible carbohydrates’, Cell Metabolism. Elsevier Inc., 20(5), pp. 779–786. doi: 10.1016/j.cmet.2014.07.003.

Soper, G. A. (1939) ‘The Curious Case of Typhoid Mary’, Bulletins of the NY Academy of Medicine, 15(10), pp. 698–712. Available at: http://www.ncbi.nlm.nih.gov/pmc/articles/PMC1911442/.

Sorbara, M. T. and Pamer, E. G. (2019) ‘Interbacterial mechanisms of colonization resistance and the strategies pathogens use to overcome them’, Mucosal immunology. NIH Public Access, 12(1), p. 1. doi: 10.1038/S41385-018-0053-0.

Stecher, B. et al. (2005) ‘Comparison of Salmonella enterica serovar typhimurium colitis in germfree mice and mice pretreated with streptomycin’, Infection and Immunity. American Society for Microbiology, 73(6), pp. 3228–3241. doi: 10.1128/IAI.73.6.3228-3241.2005/ASSET/1DE36183-8D6A-4505-B96E-D262400DE619/ASSETS/GRAPHIC/ZII0060548710007.JPEG.

Suwandi, A. et al. (2019) ‘Std fimbriae-fucose interaction increases Salmonella-induced intestinal inflammation and prolongs colonization’, PLOS Pathogens. Public Library of Science, 15(7), p. e1007915. doi: 10.1371/JOURNAL.PPAT.1007915.

Thiennimitr, P. et al. (2011) ‘Intestinal inflammation allows Salmonella to use ethanolamine to compete with the microbiota’, Proceedings of the National Academy of Sciences of the United States of America, 108(42), pp. 17480–17485. doi: 10.1073/PNAS.1107857108/SUPPL_FILE/PNAS.201107857SI.PDF.

Wardman, J. F. et al. (2022a) ‘Carbohydrate-active enzymes (CAZymes) in the gut microbiome’, Nature Reviews Microbiology. Springer Science and Business Media LLC. doi: 10.1038/s41579-022-00712-1.

Wardman, J. F. et al. (2022b) ‘Carbohydrate-active enzymes (CAZymes) in the gut microbiome’, Nature Reviews Microbiology. Springer US, 0123456789. doi: 10.1038/s41579-022-00712-1.

