## Supplemental figures for "Multi-omics reveal *Salmonella*-liberated dietary L-arabinose promotes expansion in superspreaders"

#### Supplemental figure 1

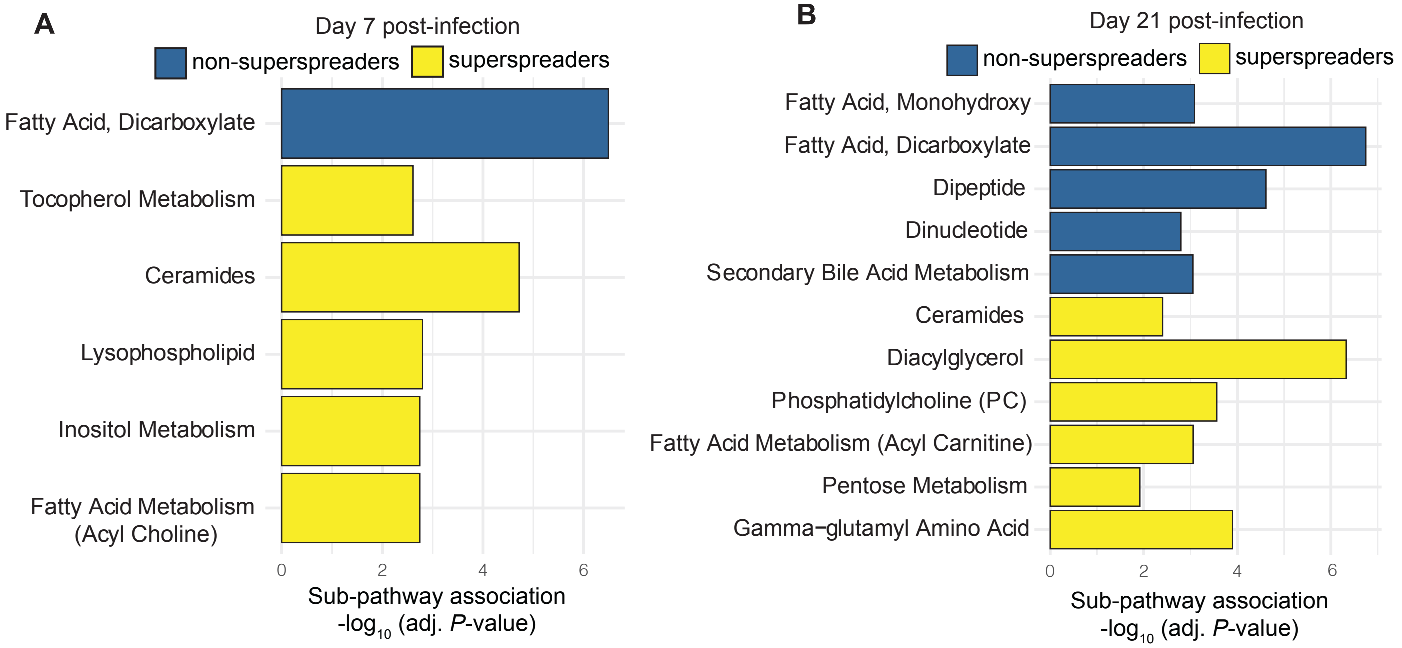

#### Supplemental figure 2

A

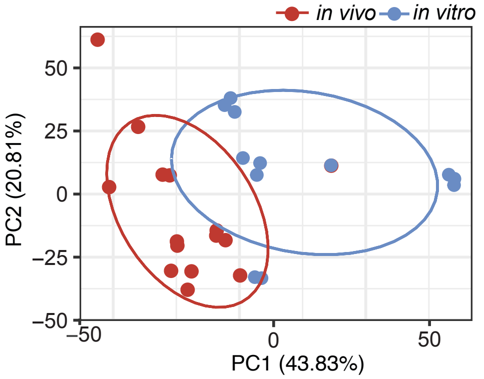


B

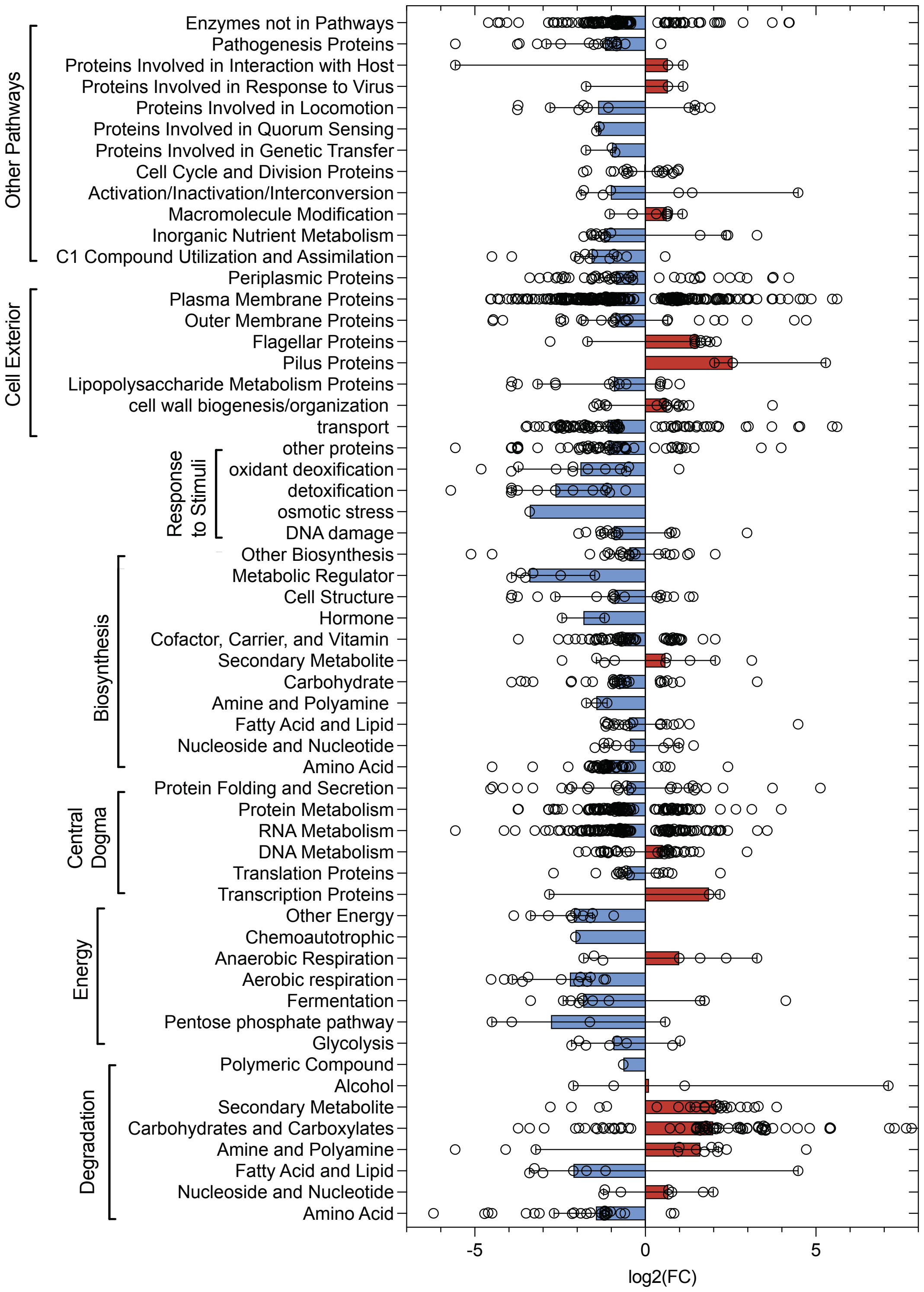

#### Supplemental figure 3

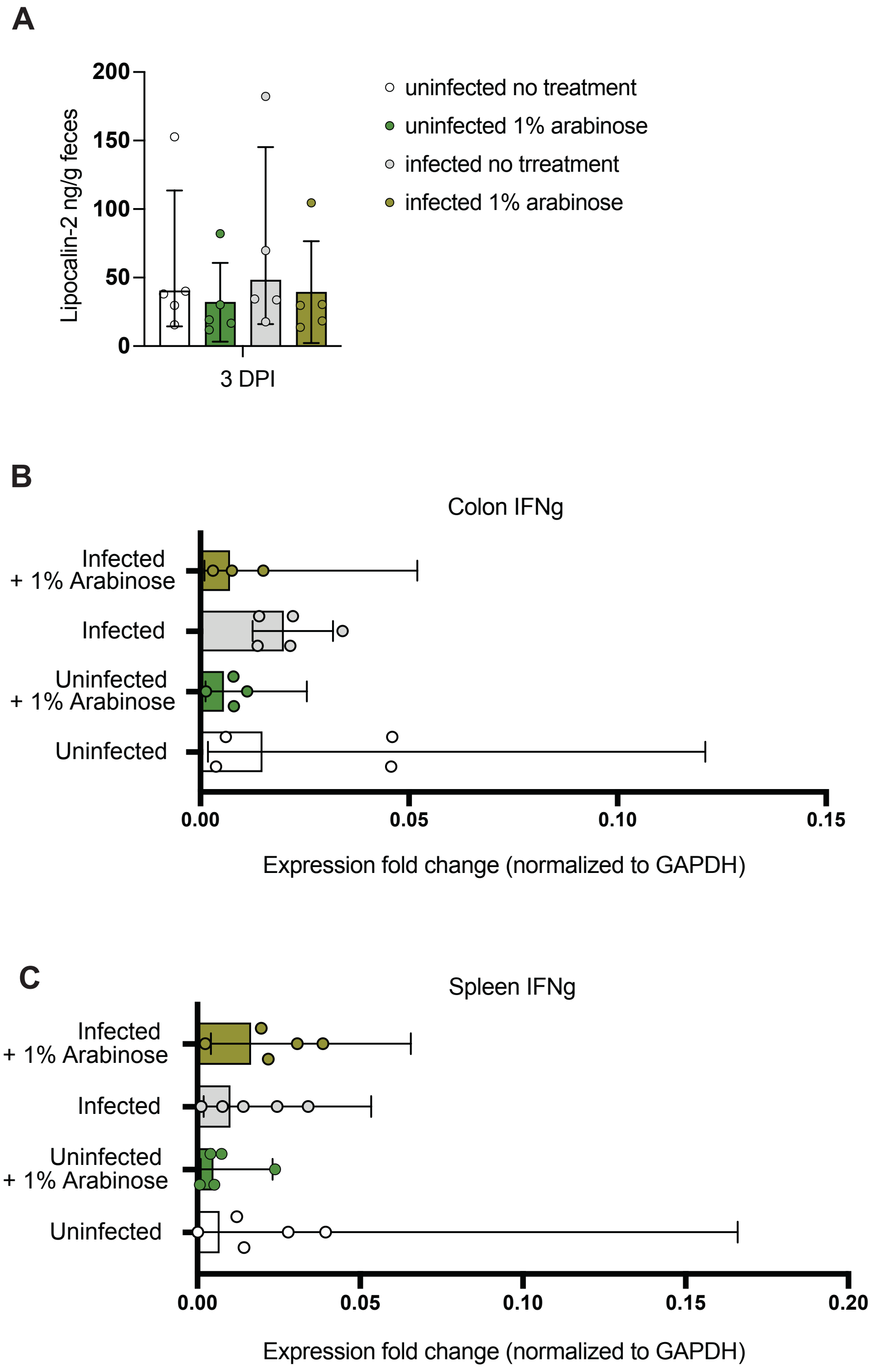

#### Supplemental figure 4

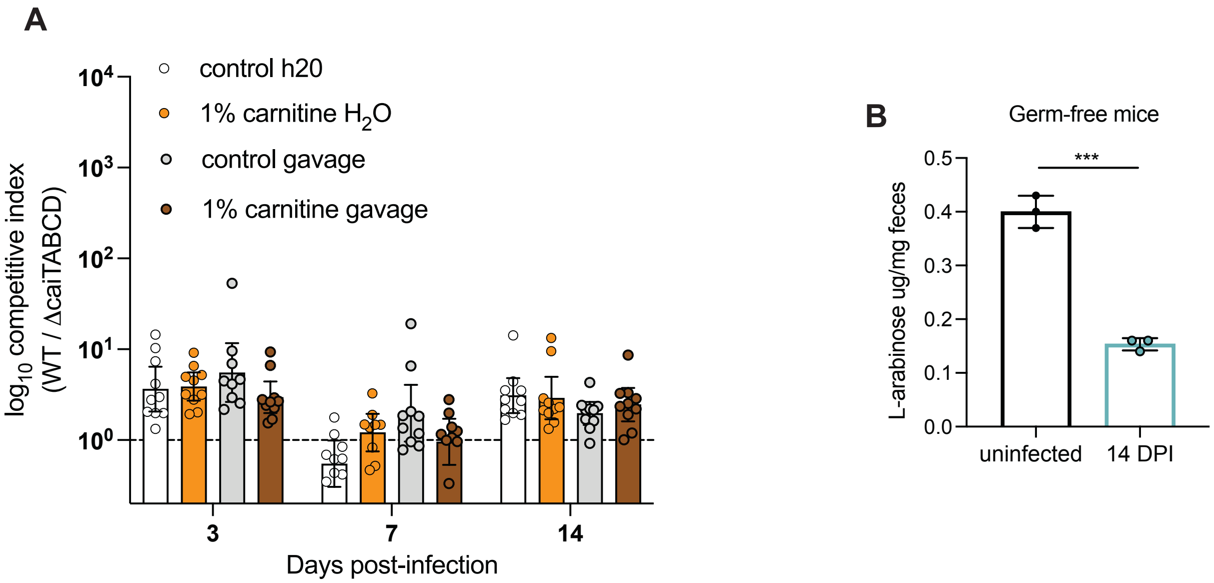

### Supplemental figure 5

## A

```
1          11         21         31         41        50
|          |          |          |          |         |
MTTSLNSRRL WLHRLCALLL GTGSAL---- ------VQAE NPIFTDVFTA   40 C. japonicus
MRKKC-SVCL WILVLLLSCL SGKSAYAATS TTIAKHIGNS NPLIDHHLGA   49 B. subtilis
M-RRL-TVRL FTAVLAALAL LTMGTPAHAT APASPSVTFT NPLAEKR--A   46 S. avermitilis
M--------- ---------- ---------- ------ANWP NPFIEQR--A   13 S. Typhimurium

DPAALVH-KG RVYLYAGRDE APDNT----- --TFFVMNEW LVYSSDDMAN   82 C. japonicus
DPVALTY-NG RVYIYMSSDD YEYNSNGTIK DNSFANLNRV FVISSADMVN   98 B. subtilis
DPHIFKHTDG Y-YYFTATVP -EYDRIVLRR ATTLQGLA-- ---TAPETTI   89 S. avermitilis
DPFILR--DG SDYYFIASVP -EYDRLEIRR ANSLEGLR-- ---AADPVVV   55 S. Typhimurium

WEAHGPGLR- -AKDFTWAKG DA------WA SQVI--ERNG --KFYWYVTV  120 C. japonicus
WTDHGAIPVA GANGANGGRG IAKWAGASWA PSIAVKKING KDKFFLYFA-  147 B. subtilis
WTKHASGVM- ---------- ----GAHIWA PEIH--FIDG --KWYVYFAA  120 S. avermitilis
WRKPESGPM- ---------- ----SQLIWA PEMH--RING --KWYIYFAA   86 S. Typhimurium

RHDD------ --TKPGFAIG VAVGDSPIGP FKDALGKALI TNDMTTDTPI  162 C. japonicus
---------- ---NSGGGIG VLTADSPIGP WTDPIGKPLV TPSTPGMSGV  184 B. subtilis
GSTS-----D VWAIRMYVLE SGAANPLTGS W--------- TEKGQIATPV  156 S. avermitilis
THTQALDKLG MFQHRMFALE CADADPLTGK W--------- TEKGQIKTPF  127 S. Typhimurium

DWDDIDPSVF IDDDGQAYLF WGNTRP---- ---------- -----RYAKL  193 C. japonicus
VW-LFDPAVF VDDDGTGYLY AGGGVPGVSN PTQ---GQWA NPKTARVIKL  230 B. subtilis
SSFSLDATTF VVN-GVRHLA WAQRNPAEDN NTSLFIAKMA NPWT------  199 S. avermitilis
DTFALDATTF YHQ-GKQWYL WAQKAPDIAG NSNIYLAELE NPWT------  170 S. Typhimurium

KKNMVELDGP IRAIEGLPEF T--------- EAIWVHKYQD NYYLSY----  230 C. japonicus
GPDMTSVVGS ASTIDA-PFM F--------- EDSGLHKYNG TYYYSY----  266 B. subtilis
------ISGT PTEISQ-PTL SWETVGYKVN EGPAVIQHGG KVFLTYSASA  242 S. avermitilis
------IKGE PVRLSK-PEY DWECRGFWVN EGPAVVVHGD KLFISYSASA  213 S. Typhimurium
```
