## Supplementary material for "Multi-omics reveal *Salmonella*-liberated dietary L-arabinose promotes expansion in superspreaders": Key resources table

| REAGENT or RESOURCE | SOURCE | IDENTIFIER |
|---|---|---|
| **Bacterial and virus strains** | | |
| *Salmonella enterica* serovar Typhimurium strain SL1344 | Monack lab strain collection | N/A |
| *S.* Typhimurium Δ*orgA*Δ*ssaV:: Kan*$^R$ | Monack lab strain collection | N/A |
| *S.* Typhimurium Δ*caiTABCD:: Kan*$^R$ | This paper | N/A |
| *S.* Typhimurium Δ*araBAD:: Kan*$^R$ | Monack lab strain collection | N/A |
| *S.* Typhimurium Δ*orgA*Δ*ssaV*Δ*araBAD:: Kan*$^R$ | This paper | N/A |
| *S.* Typhimurium Δ*0148:: Kan*$^R$ | This paper | N/A |
| *S.* Typhimurium Δ*orgA*Δ*ssaV*Δ*0148:: Kan*$^R$ | This paper | N/A |
| **Chemicals, peptides, and recombinant proteins** | | |
| LB Agar, Miller | Fischer Scientific | Cat# BP9724-2 |
| LB Broth, Miller | Fischer Scientific | Cat# BP1426-500 |
| Carbenicillin disodium salt | GoldBio | Cat# C-103-5 |
| Penicillin- streptomycin | Thermo Fisher Scientific | Cat# 15-140-122 |
| Kanamycin monosulfate | GoldBio | Cat# K-120-5 |
| Chloramphenicol | GoldBio | Cat# C-105-5 |
| L-(+)-Arabinose | Millipore Sigma | Cat# C973P47 |
| NdeI | New England Bioloabs | Cat# R0111S |
| XhoI | New England Biolabs | Cat# R0146S |
| **Critical commercial assays** | | |
| Phusion® High-Fidelity DNA Polymerase | New England BioLabs | Cat# M0530 |
| MinElute PCR Purification Kit | Qiagen | Cat# 28004 |
| RNeasy PowerMicrobiome Kit | Qiagen | Cat# 26000-50 |
| Rneasy Mini Kit | Qiagen | Cat# 74106 |
| Illumina® Stranded Total RNA Prep, Ligation with Ribo-Zero Plus | Illumina | Cat# 20040525 |
| IDT® for Illumina® RNA UDIndexes Set A, Ligation | Illumina | Cat# 20040553 |
| SuperScript III First-Strand Synthesis Kit | Thermo Fisher Scientific | Cat #18080051 |
| FastStart Universal SYBR Green Master Mix | Sigma Aldrich | Cat #4913850001 |
| Lipocalin-2 (LCN2) Mouse ELISA Kit | Invitrogen | **Cat# EMLCN2** |
| Qubit™ RNA High Sensitivity (HS), Broad Range (BR), and Extended Range (XR) Assay Kits | Thermo Fisher Scientific | Cat# Q10210 |
| **Experimental models: Organisms/strains** | | |
| 129SvJ/X1 | Jackson Laboratory | Cat# 000691 |
| GF Swiss-Webster | Sonnenburg Lab | N/A |
| **Rodent diet** | | |
| Standard diet | Envigo-Teklad | TD.2018 |
| Polysaccharide-defined (arabinose-free) diet | Envigo-Teklad | TD.150689 |

| | | |
|---|---|---|
| **Oligonucleotides** | | |
| Please see Table S2. | This study | |
| **Recombinant DNA** | | |
| pKD4 | Monack lab plasmid collection | N/A |
| pKD46 | Monack lab plasmid collection | N/A |
| pCP20 | Monack lab plasmid collection | N/A |
| **Software and algorithms** | | |
| GraphPad Prism version 8.4.3 for MacOS | GraphPad Software Inc | https://www.graphpad.com/scientific-software/prism/ |
| R(v4.0.2) | R Core Team | https://www.r-project.org/ |
| RStudio(v1.3) | RStudio Team | https://www.rstudio.com/ |
| tidyverse(v1.3.0) | Wickham et al., 2019 | https://www.tidyverse.org/ |
| ggplot2(v3.3.2) | Wickham, 2016 | https://cran.r-project.org/web/packages/ggplot2/index.html |
| Nfcore/rnaseq pipeline development branch | (Ewels et al., 2020) | https://nf-co.re/rnaseq |
| DESeq2 v1.28.0 | (Love et al., 2014) | https://bioconductor.org/packages/release/bioc/html/DESeq2.html |
| XNomial v1.0.4 | William R. Engels, University of Wisconsin, Madison – Genetics Department | https://cran.r-project.org/web/packages/XNomial/vignettes/XNomial.html#use |
