## Supplemental tables for "Multi-omics reveal *Salmonella*-liberated dietary L-arabinose promotes expansion in superspreaders"

**Table S1: Related to Figure 5. SL1344 predicted glycoside hydrolases**

| Gene ID | Cazyme family | | Reference accession |
|---|---|---|---|
| SL1344_0018 | GH18 | | CBW16119.1 |
| SL1344_3087 | GH23 | | CBW19186.1 |
| SL1344_0016 | GH108 | | CBW16117.1 |
| SL1344_0042 | GH31 | | CBW16143.1 |
| SL1344_0148 | GH43 | | CAB89837.1 |
| SL1344_0234 | GH19 | | CBW16336.1 |
| SL1344_0255 (MltD) | GH23 | | CBW16357.1 |
| SL1344_0396 | GH13 | | CBW16496.1 |
| SL1344_0966 (NucD) | GH24 | | CBW17063.1 |
| SL1344_1119 (FlgJ) | GH73 | | CBW17215.1 |
| SL1344_1146 (NagZ) | GH3 | | CBW17242.1 |
| SL1344_1188 | GH33 | | CBW17283.1 |
| SL1344_1251 (CelF) | GH4 | | CBW17347.1 |
| SL1344_1488 (GlgX) | GH13 | | CBW17583.1 |
| SL1344_1489 | GH13 | | CBW17584.1 |
| SL1344_1490 | GH13 | | CBW17585.1 |
| SL1344_1724 (TreA) | GH37 | | CBW17819.1 |
| SL1344_1727 (MltE) | GH23 | | CBW17822.1 |
| SL1344_1846 | GH105 | | CBW17940.1 |
| SL1344_1892 (AmyA) | GH13 | | CBW17987.1 |
| SL1344_1956 | GH19 | | CBW18053.1 |
| SL1344_2144 (BglX) | GH3 | | CBW18240.1 |
| SL1344_2529 (YfhD) | GH23 | | CBW18629.1 |
| SL1344_2576 | GH24 | | CBW18677.1 |
| SL1344_2686 (Nucd2) | GH24 | | CBW18788.1 |
| SL1344_2811 (MltB) | GH103 | | CBW18909.1 |
| SL1344_2857 (IagB) | GH23 | | CBW18955.1 |
| SL1344_2968 (MltA) | GH102 | | CBW19067.1 |
| SL1344_3027 (BglA) | GH1 | | CBW19126.1 |
| SL1344_3480 (MalQ) | GH77 | | CBW19575.1 |
| SL1344_3504 (GlgX) | GH13 | | CBW19598.1 |
| SL1344_3505 (GlgB) | GH13 | | CBW19599.1 |
| SL1344_3568 (TreF) | GH37 | | CBW19662.1 |
| SL1344_3570 (fragment) | GH24 | | CBW19664.1 |
| SL1344_3582 (YhjM) | GH8 | | CBW19676.1 |
| SL1344_3628 (BaX) | GH73 | | CBW19721.1 |
| SL1344_3629 (MalS) | GH13 | | CBW19722.1 |
| SL1344_3644 | GH127 | | CBW19737.1 |
| SL1344_3715 | GH31 | | CBW19809.1 |
| SL1344_3740 | GH1 | | CBW19832.1 |

| | | | |
|---|---|---|---|
| SL1344_3767 | GHnc | | CBW19857.1 |
| SL1344_3965 | GH31 | | CBW20055.1 |
| SL1344_4153 | GH23 | | CBW20241.1 |
| SL1344_4235 | GH4 | | CBW20321.1 |
| SL1344_4358 | GH30 | | CBW20445.1 |
| SL1344_4384 (TreC) | GH13 | | CBW20472.1 |
| SL1344_4509 (Slt) | GH23 | | CBW20598.1 |
| SL1344_P2_0087 (PilT) | GH23 | | CCF76895.1 |

**Table S2: Related to STAR Methods. Oligonucleotides used in this study**

| Primer name: description | Sequence 5'-3' | Source |
|---|---|---|
| SR114: caiTABCD forward lambda | AAATCGGGAATTGAACCGAAGGTTTTTTTCCG CCATTAAGTGTAGGCTGGAGCTGCTTC | IDT |
| SR115: caiTABCD reverse lambda | CGGATCGCGTTTTTCGGCAAATGCCTGCGGTC CTTCGAGCCATATGAATATCCTCCTTAG | IDT |
| SR116: caiTABCD forward verification | CTGGCTCAACAATATTGAACGC | IDT |
| SR117: caiTABCD reverse verification | GCGAGTGGGCCAATATAAACAC | IDT |
| SR118: caiTABCD reverse/internal verification | CGCGATGGTGTTATACGCC | IDT |
| SR119: araBAD forward verification | GACCAGGACGACAGAGCTTCC | IDT |
| SR120: araBAD reverse verification | CAGATTCATCAACGCGCCC | IDT |
| SR121: araBAD reverse-internal verification | GCGTCAGGGTATAGCTGCTTTCATACTC | IDT |
| SR148: 0148 lamda forward | AAACCCGTTTATTGAACAACGTGCCGATCCGTT TATTTTAGTGTAGGCTGGAGCTGCTTC | IDT |
| SR149: 0148 lamda reverse | TGGCGGCACGCCAAAATCAGGCATCCCGTTTT CGTCCCAGCATATGAATATCCTCCTTAG | IDT |
| SR150: 0148 forward verification primer | CGGCGTTGGCTATCTGATTA | IDT |
| SR151: 0148 reverse verification primer | CAATATCAGGTGCTCACACGTCTG | IDT |
| SR150: 0148 reverse-internal verification primer | CTGGGAATGTTCCAGCATCG | IDT |
| Mouse IFNγ qPCR forward | 5'-AGCGGCTGACTGAACTCAGATTGTAG-3' | IDT |
| Mouse IFNγ qPCR reverse | 5'-GTCACAGTTTTCAGCTGTATAGGG-3' | IDT |
